# H3K9me1/2 methylation limits the lifespan of *C. elegans*

**DOI:** 10.1101/2021.10.27.466082

**Authors:** Meng Huang, Minjie Hong, Chengming Zhu, Di Chen, Xiangyang Chen, Shouhong Guang, Xuezhu Feng

## Abstract

Histone methylation plays crucial roles in the development, gene regulation and maintenance of stem cell pluripotency in mammals. Recent work shows that histone methylation is associated with aging, yet the underlying mechanism remains unclear. In this work, we identified a class of histone 3 lysine 9 mono-/dimethyltransferase genes (*met-2, set-6, set-19, set-20, set-21, set-32* and *set-33*), mutations in which induce synergistic lifespan extension in the long-lived DAF-2 (IGF-1 receptor) mutant in *C. elegans*. These histone methyltransferase plus *daf-2* double mutants not only exhibited an average lifespan nearly three times that of wild-type animals and a maximal lifespan of approximately 100 days, but also significantly increased resistance to oxidative and heat stress. Synergistic lifespan extension depends on the transcription factor DAF-16 (FOXO). mRNA-seq experiments revealed that the mRNA levels of class I DAF-16 target genes, which are activated by DAF-16, were further elevated in the double mutants. Among these genes, *F35E8.7, nhr-62, sod-3, asm-2* and *Y39G8B.7* are required for the lifespan extension of the *daf-2;set-21* double mutant. In addition, treating *daf-2* animals with the H3K9me1/2 methyltransferase G9a inhibitor also extends lifespan and increases stress resistance. Therefore, investigation of DAF-2 and H3K9me1/2 methyltransferase deficiency-mediated synergistic longevity will contribute to a better understanding of the molecular mechanisms of aging and therapeutic applications.

## Introduction

Lifespan is governed by complex interactions between genetic and environmental factors. The perturbation of insulin/insulin-like signaling (IIS), target of rapamycin (TOR) pathway, and mitochondrial functions have been shown to extensively modulate lifespan and health (1–3). These genetic manipulations often lead to significant changes in gene expression at both the transcriptional and translational levels. Inhibition of DAF-2, the *C. elegans* ortholog of the insulin growth factor 1 (IGF-1) receptor, doubles adult lifespan by activating the DAF-16 (FOXO) transcription factor to regulate downstream genes involved in stress resistance, detoxification, and metabolism (4–9).

In addition to genetic regulation, aging is also modulated by epigenetic processes. Epigenetic marks, including histone acetylation and methylation, as well as the associated chromatin states, are altered during aging. Age-dependent loss of chromatin repression is correlated with alterations in gene expression patterns, which have been documented in species ranging from *C. elegans* to humans (10–13). For example, the methylation of histone 3 lysine 4 methylation (H3K4me) is one of the posttranslational histone modifications that marks on regions of active transcription (14). H3K4me is deposited by the MLL/COMPASS complex, which travels with elongating RNA Polymerase II during transcription (15). In *C. elegans*, animals with reductions in COMPASS complex subunits (*wdr-5, ash-2,* and *set-2*) live longer than wild-type individuals (16).

However, little is known about the importance of repressive histone modification for aging marks. The H3K27me3 demethylase UTX-1 regulates lifespan independently of the presence of the germline but in a manner that depends on the insulin-FOXO signaling pathway (17). MET-2, a mammalian H3K9 methyltransferase SETDB1 homolog (18), monomethylates and dimethylates H3K9 in *C. elegans* (19). MET-2 is necessary both for a normal lifespan (20) and for the lifespan extension of *wdr-5* mutants (21). SET-6 is a putative H3K9me2/3 methyltransferase but not H3K9me1. Although the depletion of SET-6 does not significantly increase the lifespan of *C. elegans*, it may increase healthy aging (22). SET-25 is a tri-methylase for H3K9 (19, 23). However, the deletion of SET-25 does not significantly change the worm lifespan (24).

The enzymes responsible for histone lysine methylation are called histone methyltransferases (HMTs). HMTs typically contain a conserved catalytic domain called SET, which stems from **S**u(var)3–9, **E**nhancer of zeste, and **T**rithorax, the first HMTs known to carry this domain (25). The *C. elegans* genome encodes 38 SET domain-containing proteins, of which five are essential for viability (26, 27). However, the biological roles of most SET proteins are largely unknown.

To investigate the function of repressive histone methylation in lifespan regulation, we selected a number of putative H3K9 methyltransferases and tested whether the loss-of-function of these SET proteins could change the lifespan of *C. elegans*. Interestingly, we found that the H3K9me1/2, but not H3K9me3, mutants exhibited a synergistic lifespan extension with *daf-2* mutation. These animals show an average lifespan of approximately sixty days, which is approximately 70% longer than that of *daf-2* worms and is three times as long as that of wild-type N2 animals. The double mutants exhibited a maximal lifespan of approximately 100 days. The synergistic lifespan extension of DAF-2 and lifespan-limiting histone methyltransferase mutants depend on the transcription factor DAF-16 (FOXO). mRNA-seq experiments showed that class I DAF-16 targets (genes that are activated by DAF-16) are activated in long-lived worms. Therefore, we conclude that H3K9me1/2 may limit nematode lifespan by repressing the expression of class I DAF-16 targets.

## Results

### The depletion of *set-21* extends lifespan and enhances stress resistance

To investigate the function of repressive histone methylation in lifespan regulation, we selected ten putative H3K9 methyltransferases which contain conserved catalytic SET domains (Fig. 1A). Most of these genes have unknown functions. We first generated a number of deletion alleles of *set-21* by CRISPR/Cas9 technology (Fig. S1A). The deletion of *set-21* did not significantly change the lifespan in N2 background (Fig. S1B) and subtly altered the brood size (Fig. S1C). Strikingly, knocking out *set-21* significantly extended lifespan in *daf-2(e1370)* mutant worms (Fig. 1B). The average lifespan of *daf-2(e1370);set-21(ust68)* were 59% longer than that of *daf-2(e1370)* animals (Fig. 1B). And the maximal life span of *daf-2;set-21* animals achieved approximately 90 days, which is three time as long as that of N2 animals.

**Figure 1.**
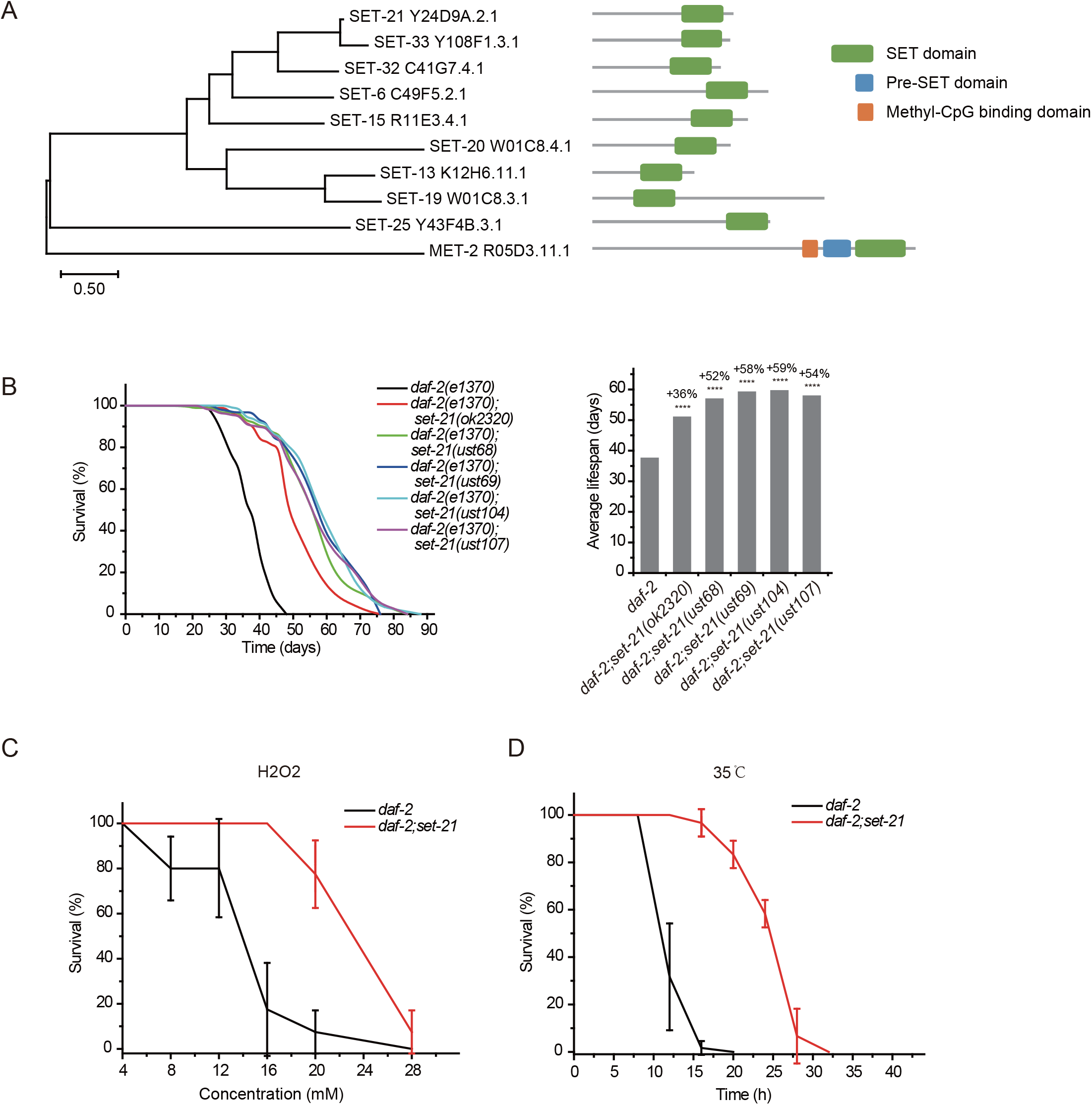
Synergistic lifespan extension and stress resistance in *daf-2;set-21* mutants. (a) Phylogenetic tree comparing the protein sequences of SET proteins that are predicted to encode H3K9 methyltransferases in *C. elegans*. (b) (Left) Survival curves and (right) average lifespan of the indicated animals. The percentage of change was compared to the average lifespan of *daf-2* animals. Asterisks indicate significant differences using two-tailed t tests. ****P < 0.0001. (c, d) Survival curves of the indicated animals. (c) oxidative and (d) heat stress. Data are presented as the mean ± s.e.m. of five independent experiments.

The extended lifespan of nematodes has been shown to correlate with the activation of stress response genes and increased stress resistance (8, 28, 29). *daf-2;set-21* worms also revealed a much higher resistance to oxidative stress via hydrogen peroxide treatment and heat shock stress than *daf-2* animals (Figs. 1C-D).

### The depletion of *met-2* extends lifespan in *daf-2* mutant worms and enhances stress resistance

The remarkable lifespan extension in *daf-2;set-21* animals inspired us to re-investigate the role of histone H3 lysine 9 methylation in *C. elegans*. MET-2 is the mammalian SETDB1 homolog, which is involved in mono- and dimethylation of H3K9 (Fig. 2A) (18, 19). *met-2* mutants exhibited a modest shorter lifespan than wild-type N2 animals (Fig. 2B) (20). Strikingly, *daf-2;met-2* double mutants revealed an average lifespan of approximately 47 days, which is 30% longer than that of *daf-2* mutation alone and is 2.3 times as long as that of wild-type N2 animals (Fig. 2B). The depletion of *met-2* enhanced the oxidative stress resistance and heat stress resistance in both N2 and *daf-2* mutant worms (Figs. 2C-F). These results suggest that *met-2* and H3K9me1/2 may play different roles in lifespan regulation in N2 and *daf-2* animals. Alternatively, the depletion of *daf-2* may provide a sensitized genetic background to identify genes that are involved in ageing and lifespan regulation.

**Figure 2.**
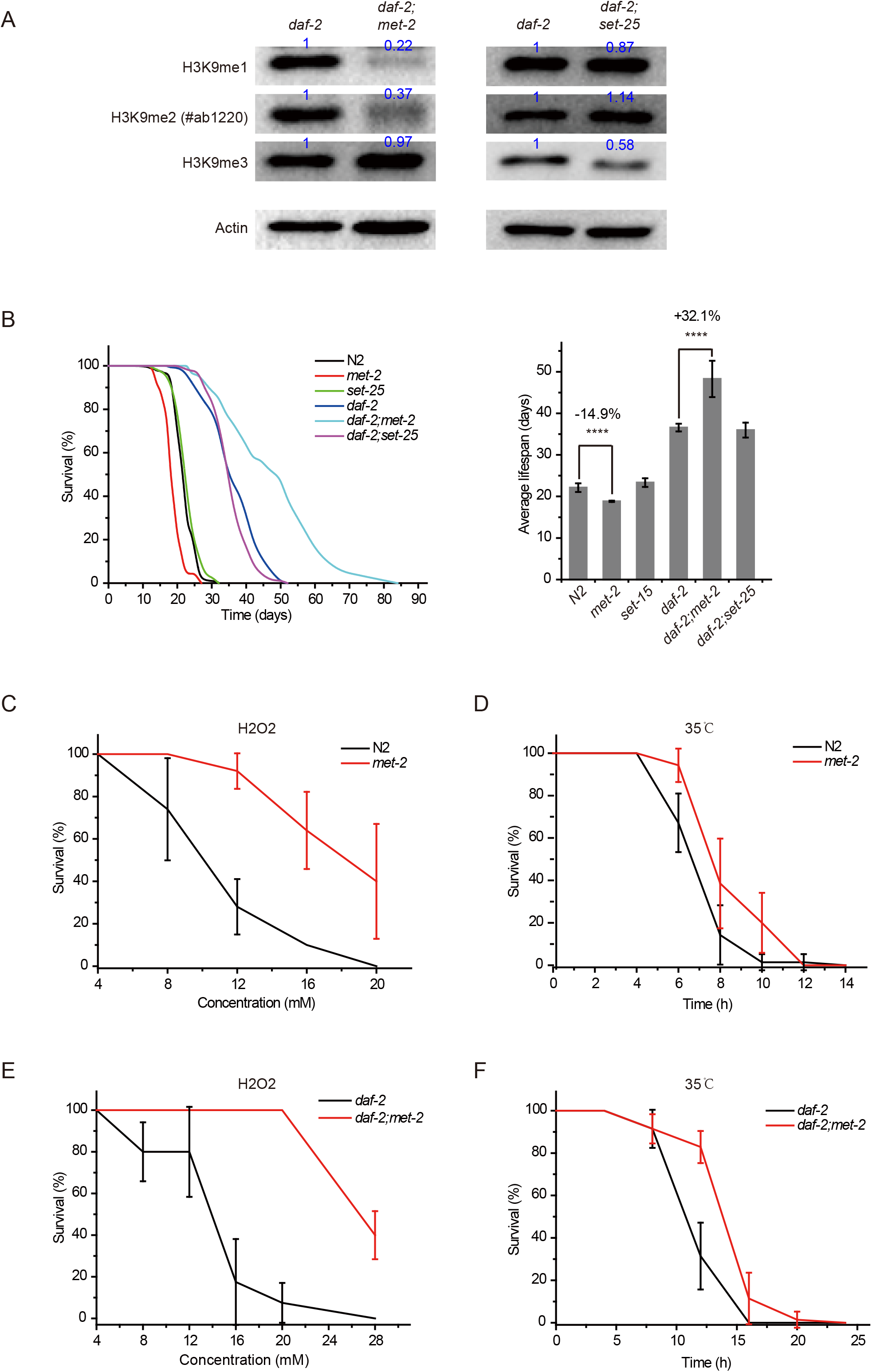
*daf-2;met-2* mutants revealed extended lifespan and increased resistance to oxidative and heat stress. (a) Western blotting of L4 stage animals with the indicated antibodies. Numbers indicate the scanned density by ImageJ. The original files of the full raw unedited blots and figures with the uncropped blots with the relevant bands clearly labelled are provided in the source data 1. (b) (Left) Survival curves of indicated animals. (Right) Histogram displaying the average lifespan of the indicated animals. mean ± s.e.m. of three independent experiments. ****P < 0.0001. (c, d, e, f) Survival curves of the indicated animals. (c, e) oxidative and (d, f) heat stress. Data are presented as the mean ± s.e.m. of five independent experiments.

SET-25 is a tri-methylase for H3K9 (Fig. 2A) (19). However, deletion of SET-25 did not significantly change the worm lifespan in either the wild-type N2 or *daf-2* background animals (Fig. 2B) (24).

### Identification of *set* genes required for lifespan limitation

Using N2 and the sensitized *daf-2* background, we tested the lifespan of the other seven putative H3K9 methyltransferase mutants. We acquired additional *set* mutants from CGC and also generated alleles by CRISPR/Cas9 technology (Fig. S2A). Among them, the deletion of *set-6, set-19, set-20* and *set-32* modestly increased the lifespan compared to that of wild-type N2 animals (Fig. 3A). However, in the *daf-2* mutant background, mutations of *set-6, set-19, set-20, set-32* and *set-33* exhibited a striking synergistic lifespan extension. While the lifespan of *daf-2;set-20* and *daf-2;set-32* are approximately 60% longer than that of *daf-2* worms, the *daf-2;set-6* and *daf-2;set-19* live 70% longer than that of *daf-2* animals (Fig. 3B). Especially, *daf-2;set-19* exhibits a maximal lifespan of approximately 100 days, which is five times as long as the average lifespan of N2 animals. *daf-2;set-6, daf-2;set-19, daf-2;set-20, daf-2;set-32, daf-2;set-33* had fewer brood sizes than the single mutants (Fig. S2B). The long-lived animals were also more resistant than control animals to the oxidative stress induced by hydrogen peroxide (Fig. 3C) and heat stress (Fig. 3D).

**Figure 3.**
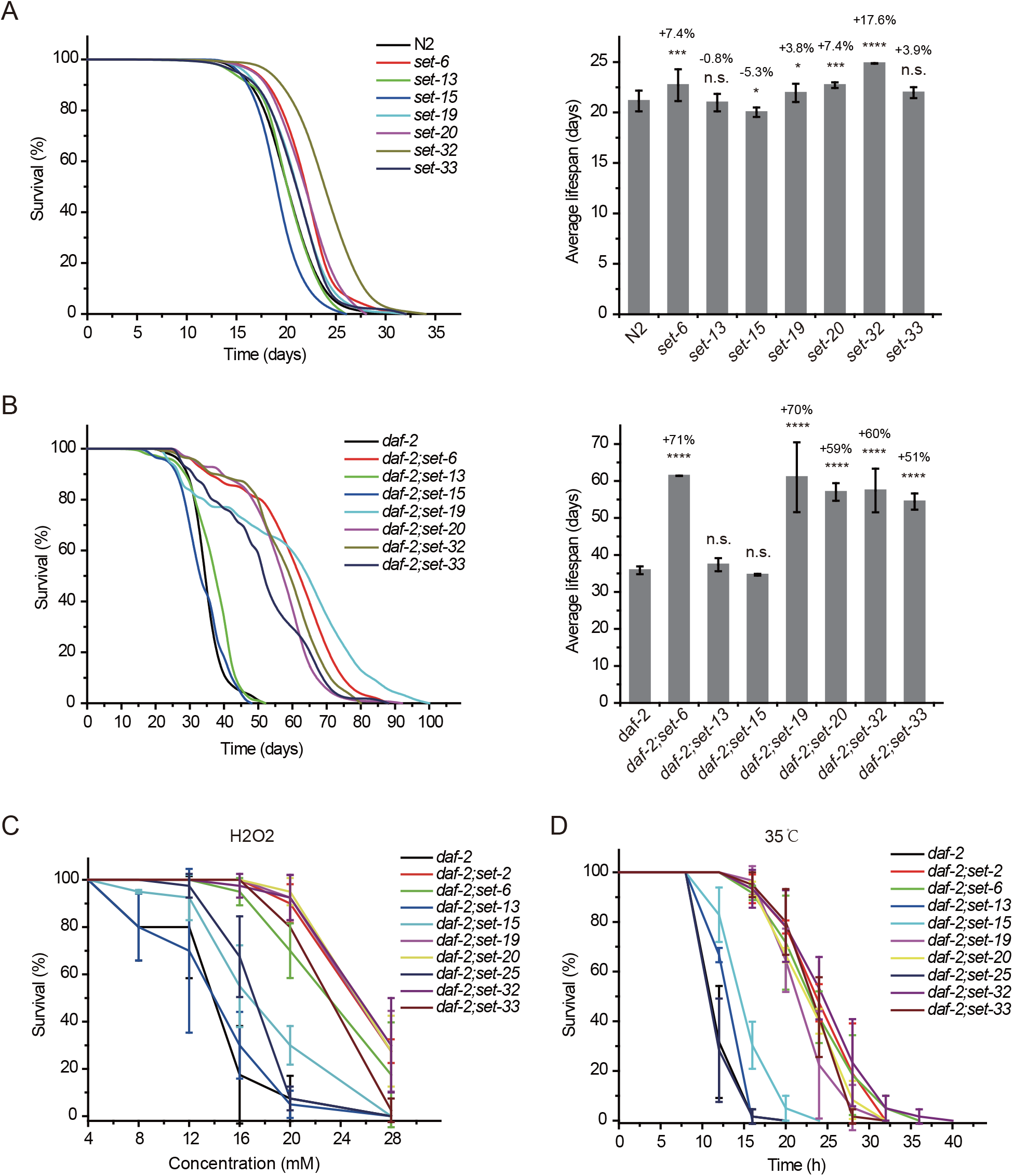
Synergistic lifespan extension of *set-6, set-19, set-20, set-32* and *set-33* with *daf-2* animals. (a) (Left) Survival curves of indicated animals. (Right) Histogram displaying the average lifespan of the indicated animals. mean ± s.e.m. of three independent experiments. The percentage of change was compared to the average lifespan of N2 animals. *P < 0.05; **P < 0.01; ***P < 0.001; ****P < 0.0001; n.s., not significant. (b) (Left) Survival curves of indicated animals. (Right) Histogram displaying the average lifespan of the indicated animals. mean ± s.e.m. of three independent experiments. The percentage of change was compared to the average lifespan of *daf-2* animals. *P < 0.05; **P < 0.01; ***P < 0.001; ****P < 0.0001; n.s., not significant. (c, d) Survival curves of the indicated animals. (c) oxidative and (d) heat stress. Data are presented as the mean ± s.e.m. of five independent experiments.

To test whether SET-6, SET-19, SET-20, SET-21, SET-32 and SET-33 act in the same genetic pathway to regulate lifespan, we crossed *set-21(ust68)* to other *set* mutants. As expected, the triple mutants *daf-2;set-21;set-6, daf-2;set-21;set-19, daf-2;set-21;set-20, daf-2;set-21;set-32,* and *daf-2;set-21;set-33* did not significantly extend the lifespan further than the double mutants (Fig. 4A). Thus, these *set* genes probably act in the same genetic pathway to regulate lifespan.

**Figure 4.**
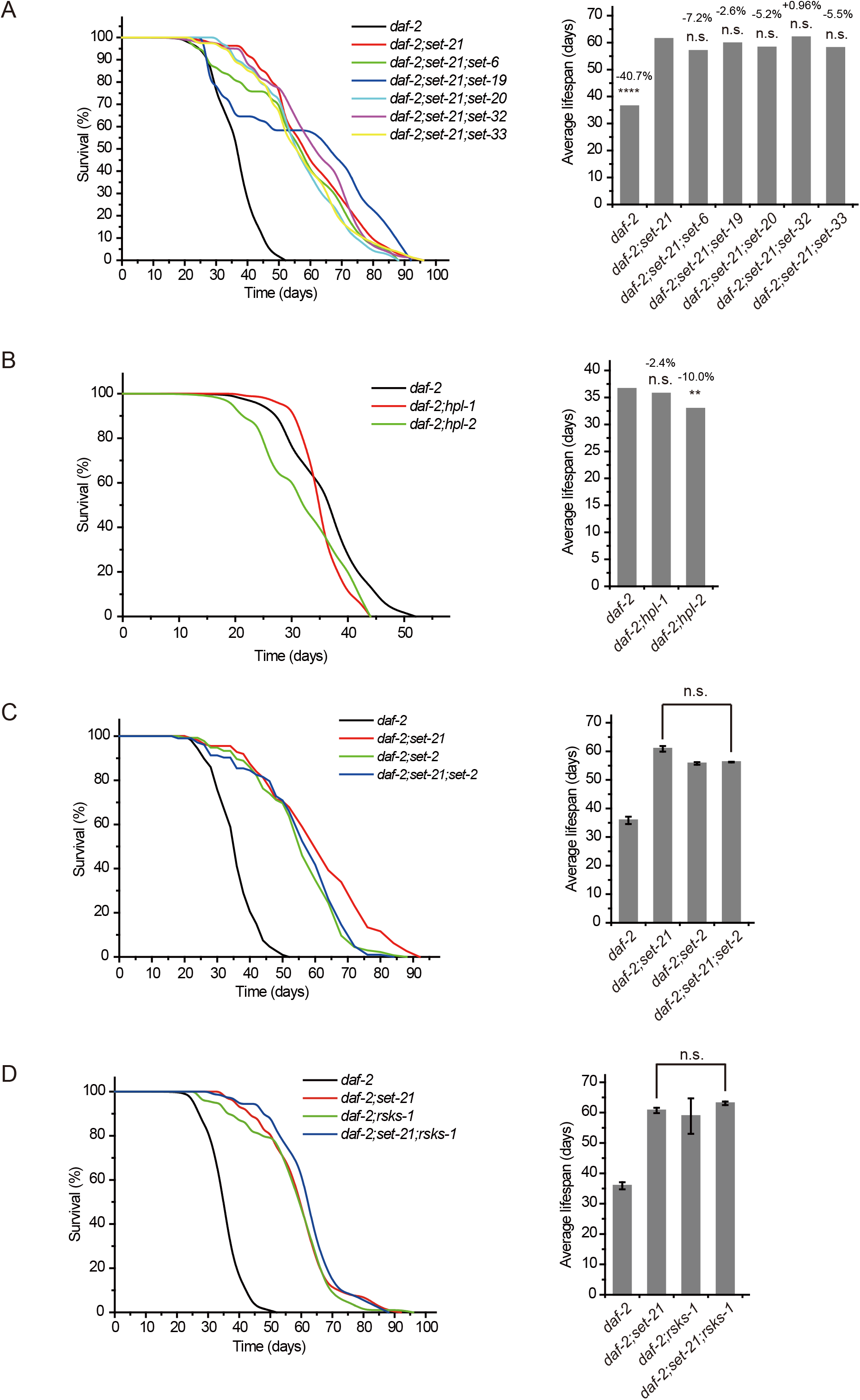
Genetic pathway analysis of *set* genes in lifespan regulation. (a) (Left) Survival curves of the indicated animals. (Right) Histogram displaying the average lifespan of the indicated animals. The percentage change was compared to the lifespan of *daf-2;set-21* animals. ****P < 0.0001; n.s., not significant. (b) (Left) Survival curves of the indicated animals. (Right) Histogram displaying the average lifespan of the indicated animals. The percentage change was compared to the lifespan of *daf-2* animals. **P < 0.01; n.s., not significant. (c, d) (Left) Survival curves of the indicated animals. (Right) Histogram displaying the average lifespan of the indicated animals. means + s.e.m. of three independent experiments. n.s., not significant.

H3K9me3 is usually considered an epigenetic hallmark of heterochromatin, which is recognized by the HP1-like proteins HPL-1 and HPL-2, orthologs of human CBX3 (chromobox 3) (19, 30, 31). Consistently, double mutants of *hpl-1* and *hpl-2* with *daf-2* did not show further lifespan extension compared to the *daf-2* mutant (Fig. 4B). SET-25 is required for H3K9me3 methylation but is not involved in lifespan limitation (Fig. 2B), further supporting that H3K9me3 may be dispensable for lifespan regulation in *C. elegans*.

Previous work has shown that knocking out the H3K4me3 methyltransferase SET-2 extended worm lifespan (16). To test whether SET-2-dependent H3K4me3 and lifespan-limiting H3K9 methyltransferases act in the same genetic pathway to regulate lifespan, we crossed *set-2* to *set-21* animals. The double mutants *daf-2;set-2* and *daf-2;set-21* and the triple mutants *daf-2;set-2;set-21* exhibited lifespan extensions similar to those of *daf-2* animals (Fig. 4C), suggesting that H3K4me3 and H3K9me1/2 may function in the same genetic pathway or regulate the same cohorts of target genes for lifespan modulation.

Inhibition of RSKS-1 (S6K), the target of rapamycin (TOR) pathways, extends lifespan in *C. elegans* (32, 33), and double mutant of *daf-2;rsks-1* leads to synergistically prolonged longevity (34). However, the triple mutants *daf-2(e1370);rsks-1(ok1255);set-21(ust68)* revealed similar lifespan as long as those of *daf-2(e1370);rsks-1(ok1255)* and *daf-2(e1370);set-21(ust68)* double mutants (Fig. 4D), suggesting that either *rsks-1* and *set-21* act in the same molecular pathway to regulate lifespan or there is likely an up ceiling of maximal life span of approximately 100 days for *C. elegans*. Interesting, a report suggested that in a mutant of *age-1*, which encodes the class-I phosphatidylinositol 3-kinase catalytic subunit (PI3K(CS)), worms can survive to a median of 145-190 days at 20 degrees and a maximal life span of approximately 260 days, with nearly 10-fold extension of both median and maximum adult lifespan relative to control animals (35).

Therefore, we concluded that a number of *set* genes limit the lifespan and stress resistance in *C. elegans*.

### Lifespan-limiting SET proteins regulate class I DAF-16 target genes

The longevity phenotype of *daf-2* animals depends on the downstream DAF-16 transcription factor (4). To test the role of DAF-16 in the *daf-2;set-21*-induced synergistic longevity, we constructed a *daf-2(e1370);daf-16(mu86);set-21(ust68)* triple mutant. The *daf-16* mutation reverted the prolonged longevity phenotype of *daf-2;set-21* to an average lifespan of 23 days, which is similar to that of *set-21* or N2 alone (Fig. 5A). In the N2 background, the *daf-16* and *daf-16;set-21* mutant worms lived shorter than wild-type N2 and *set-21* animals (Fig. 5B).

**Figure 5.**
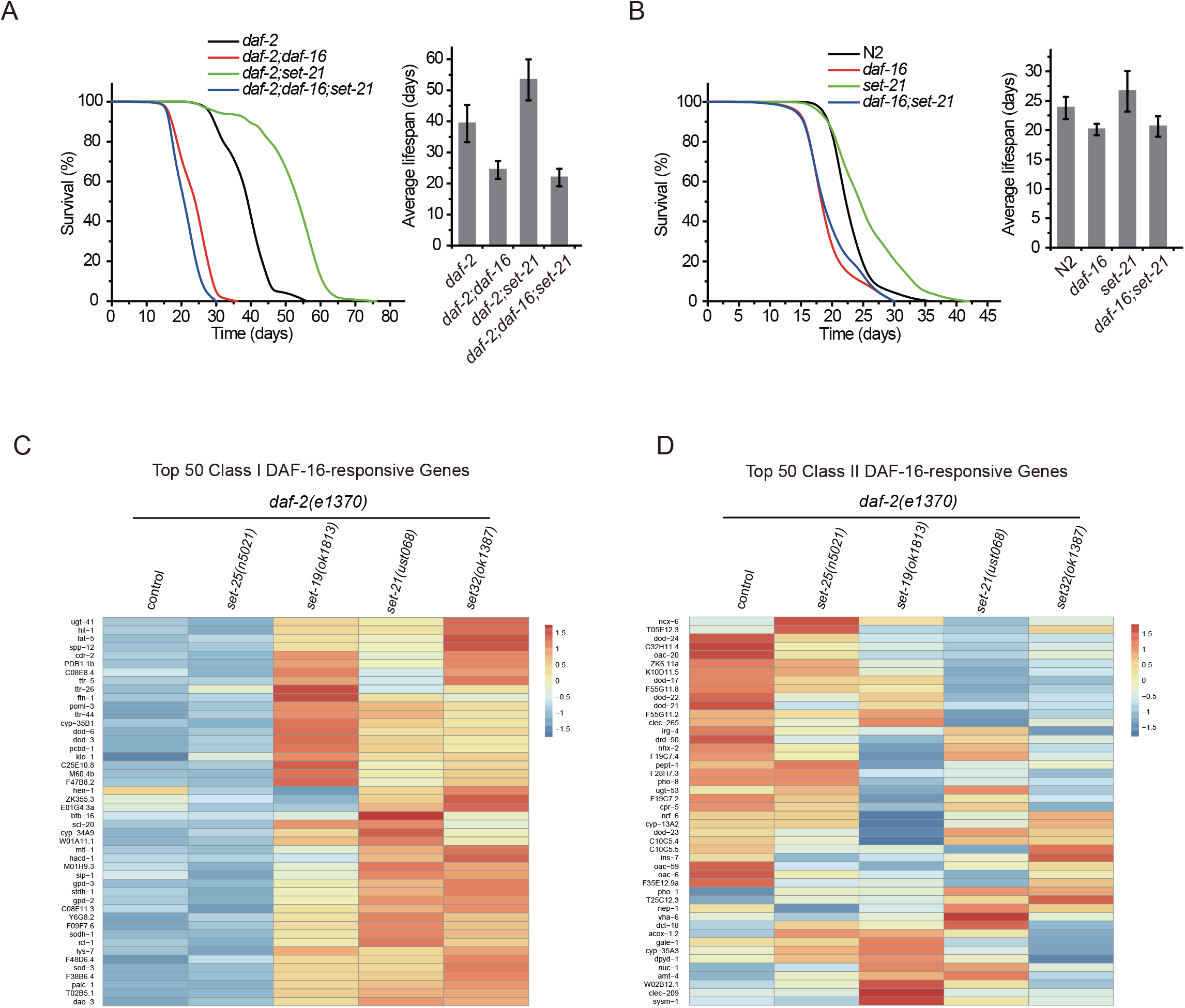
The SET proteins regulate Class I DAF-16 target genes. (a, b) (Left) Survival curves of indicated animals. (Right) Histogram displaying the average lifespan of the indicated animals. mean ± s.e.m. of three independent experiments. (c, d) Differential expression of the top 50 (c) class I and (d) class II DAF-16 target genes by mRNA-seq in the indicated animals.

DAF-16 localizes in the cytoplasm in N2 worms but accumulates in the nucleus upon *daf-2* mutation (36, 37). To determine whether the expression pattern and subcellular localization of DAF-16 are altered upon mutation of these lifespan limiting histone methyltransferases, we crossed DAF-16::GFP with *set-21*, *set-25, daf-2;set-21* and *daf-2;set-25* animals. However, the mutation of *set-21* or *set-25* did not induce a detectable change in the expression pattern and subcellular localization of DAF-16::GFP in either N2 or *daf-2* background worms (Figs. S3A-B).

Reduced insulin/IGF-1-like signaling (IIS) extends the lifespan of *C. elegans* by upregulating the stress response (class I) and downregulating other (class II) genes (38). DAF-16 directly regulates class I genes, through the DAF-16-binding element (DBE). PQM-1 is another transcriptional activator that directly controls development (class II) genes by binding to the DAF-16-associated element (DAE). We performed mRNA-seq to identify the target genes of these lifespan-limiting histone methyltransferases. Interestingly, the mRNA levels of Top 50 class I, but not class II, DAF-16 targets are all activated in long-lived *daf-2;set-19, daf-2;set-21* and *daf-2;set-32* worms, than in control animals *daf-2* and *daf-2;set-25* (Figs. 5C-D).

### Knocking out the class I DAF-16 target genes partially reverted the synergistic lifespan extension in *daf-2;set-21* animals

To confirm the change in target gene expression, we chose 10 class I DAF-16 target genes and quantified the mRNA levels by quantitative real-time PCR (qRT–PCR) in *daf-2*, *daf-2;set-21* and *daf-2;set-25* mutants (Fig. 6A). We generated deletion mutants of these genes by CRISPR/Cas9 technology (Fig. S4) and crossed the mutants into the *daf-2;set-21* background for lifespan analysis. Five of these mutants, *nhr-62, sod-3, asm-2, F35E8.7,* and *Y39G8B.7*, could partially revert the lifespan extension phenotype of *daf-2;set-21* animals (Fig. 6B). Among them, NHR-62 is a nuclear hormone receptor with DNA binding activity, which is required for dietary restriction-induced longevity in *C. elegans* (39). SOD-3 is a superoxide dismutase that is involved in the removal of superoxide radicals and required for lifespan extension in *isp-1* mutant worms (40). ASM-2 is an ortholog of human SMPD1 (sphingomyelin phosphodiesterase 1) and is involved in ceramide biosynthetic processes and sphingomyelin catabolic processes (41, 42). Strikingly, both F35E8.7 and Y39G8B.7 are predicted to encode proteins with ShK domain-like or ShKT domains, yet their functions are unknown.

**Figure 6.**
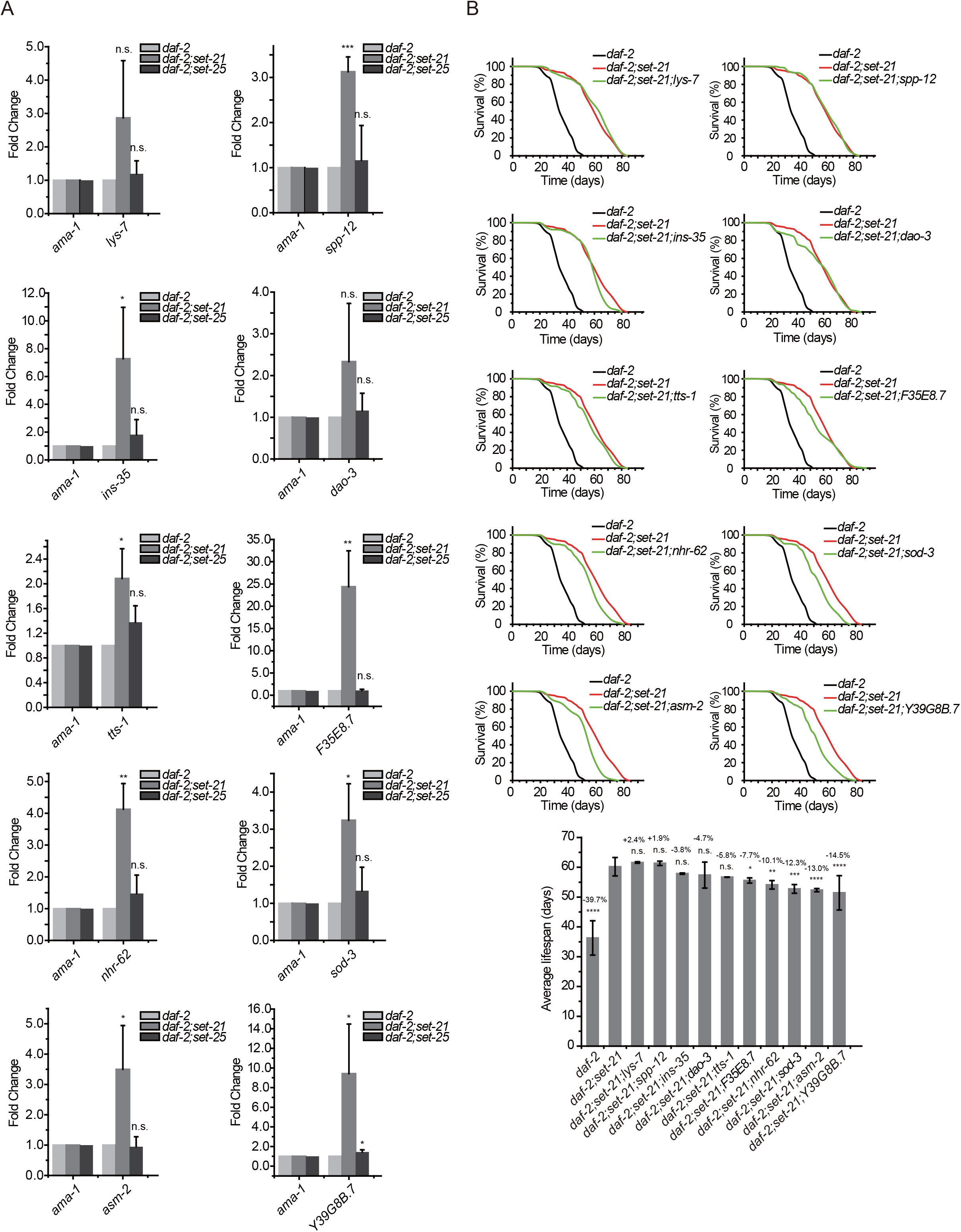
Class I DAF-16 target genes are required for the synergistic lifespan extension in *daf-2;set-21* animals. (a) Quantitative real-time PCR analysis of the indicated mRNAs. Data are presented as the mean ± s.e.m. of three independent experiments. *P < 0.05; **P < 0.01; ***P < 0.001; ****P < 0.0001; n.s., not significant. (b) (Top) Survival curves of indicated animals. (Bottom) Histogram displaying the average lifespan of the indicated animals. mean ± s.e.m. of three independent experiments. The percentage of change was compared to the average lifespan of *daf-2;set-21* mutants. *P < 0.05; **P < 0.01; ***P < 0.001; ****P < 0.0001; n.s., not significant.

Therefore, we concluded that the deletion of lifespan limiting *set* genes in *daf-2*leads to a synergistically extended lifespan by increasing DAF-16 activity.

### Lifespan-limiting SET proteins are required for H3K9me1/2 modification

To further investigate the mechanism by which these SET proteins limit *C. elegans*’ lifespan, we used Western blotting assay to examine the levels of a number of histone 3 methylation marks in L4 animals and embryos (Figs. 7A, S5A-B). Previous work have reported that SET-6 is required for H3K9me2/3 methylation (22), SET-25 is required for H3K9me3 methylation (19), and SET-32 is required for H3K23 methylation (30).

**Figure 7.**
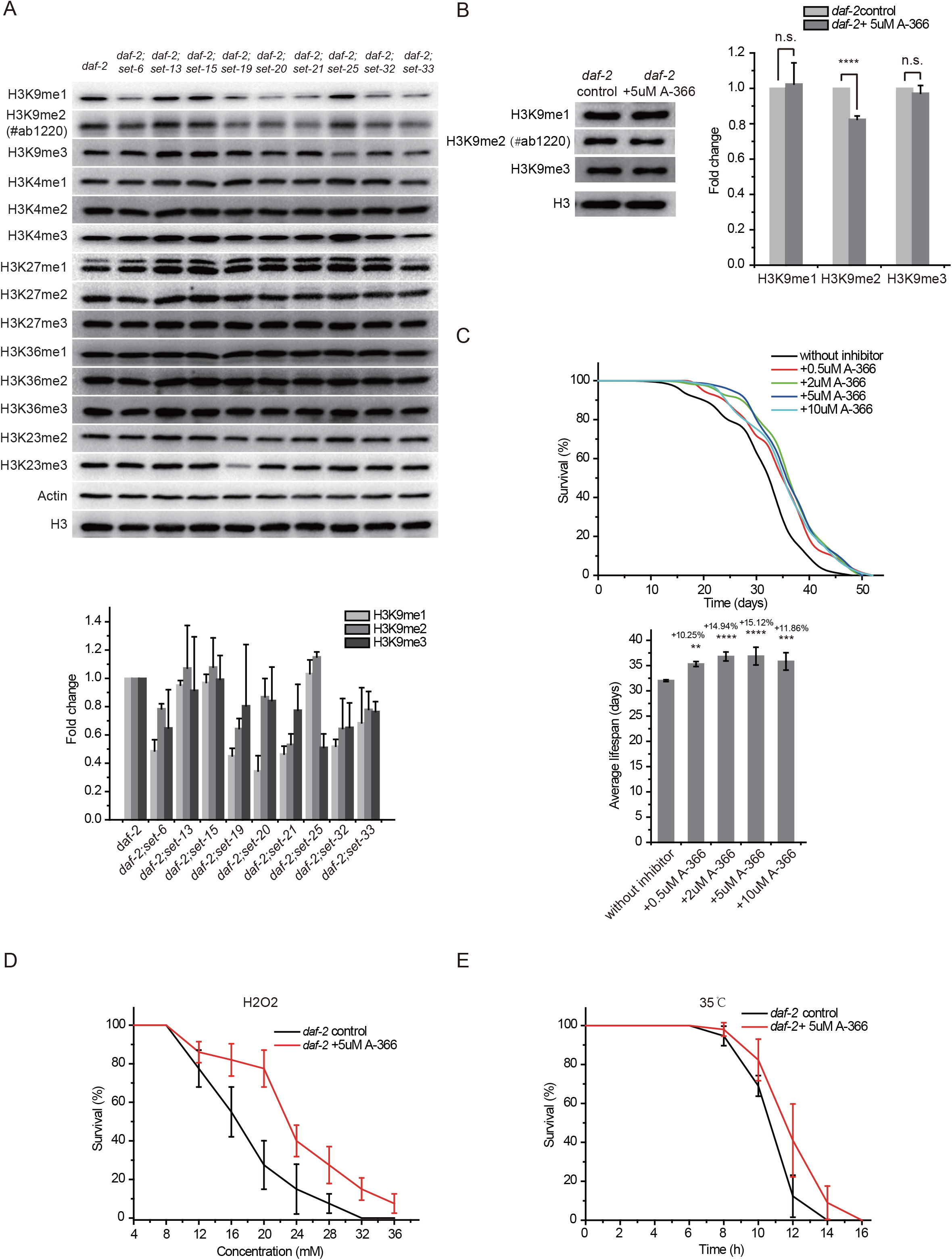
H3K9me1/2 methylation limits the lifespan and stress resistance of *daf-2* mutants. (a) (Top) Western blotting of L4 stage animals with the indicated antibodies (also see Figs. S5). (Bottom) The histogram displayed means + s.e.m. of scanned density by ImageJ from three independent experiments. The original files of the full raw unedited blots and figures with the uncropped blots with the relevant bands clearly labelled are provided in the source data 2. (b) The G9a (EHMT2) inhibitor A-366 reduced H3K9me2 levels. (Left) Western blotting of L4 stage animals with the indicated antibodies. (Right) The histogram displayed means + s.e.m. of scanned density by ImageJ from three independent experiments. ****P < 0.0001. The original files of the full raw unedited blots and figures with the uncropped blots with the relevant bands clearly labelled are provided in the source data 2. (c) (Top) Survival curves of indicated animals. (Bottom) Histogram displaying the average lifespan of the indicated animals. mean ± s.e.m. of three independent experiments. The percentage of change was compared to the average lifespan of the *daf-2* mutant. **P < 0.01; ***P < 0.001; ****P < 0.0001. (d, e) Survival curves of G9a(EHMT2) inhibitor A-366-treated *daf-2* animals upon (d) oxidative and (e) heat stress. Data are presented as the mean ± s.e.m. of five independent experiments.

In the *daf-2* mutant background, long-lived *set-6, set-19, set-20, set-21, set-32* and *set-33* mutants, but not *set-13, set-15* and *set-25* animals, decreased global H3K9me1/2 levels at the L4 larval stage (Fig. 7A). We used another anti-H3K9me2 antibody #ab176882 and confirmed that *daf-2;set-6, daf-2;set-19, daf-2;set-20, daf-2;set-21, daf-2;set-32* and *daf-2;set-33* mutants, but not *daf-2;set-13, daf-2;set-15* and *daf-2;set-25* animals, decreased global H3K9me1/2 levels at the L4 larval stage (Fig. S5A). In embryos, *set-15, set-20* and *set-32* mutants reduced H3K9me1/2 levels (Fig. S5B). The *daf-2;set-19* mutant also showed decreased H3K23me3 levels (Figs. 7A and S5B). Although H3K4me has been shown to be involved in lifespan regulation, none of the long-lived *set-6, set-19, set-20, set-21, set-32* and *set-33* mutants revealed significant changes in H3K4 methylation levels (Figs. 7A and S5B). Chromatin immunoprecipitation (ChIP) followed by qRT–PCR further revealed a modest reduction in H3K9me1/2 levels of the 10 class I DAF-16 target genes in *daf-2;set-21* mutants (Fig. S6).

Histone methyltransferase G9a, also known as euchromatic histone lysine methyltransferase 2 (EHMT2), is the human homolog of SET-6 (22) and mediates H3K9 di-methylation. Treating *daf-2* animals with the G9a inhibitor A-366 reduced H3K9me2 levels (Fig. 7B). The A-366 treatment extended the lifespan of *daf-2* worms by 15% (Fig. 7C). Moreover, A-366 also increased nematode resistance to oxidative and heat stress (Figs. 7D-E). Thus, we conclude that decreased H3K9me1/2 levels in *C. elegans* by a G9a inhibitor may increase lifespan and resistance to oxidative stress and heat stress.

## Discussion

Here, we identified a class of histone methyltransferases that limit the lifespan of *C. elegans*. These histone methyltransferases are involved in H3K9me1/2 modification and regulate Class I DAF-16 target genes. In the absence of H3K9me1/2, the binding affinity of DAF-16 to Class I target genes increases, which promotes the expression of longevity genes and anti-stress genes and extends lifespan (Fig. S7). Decreased H3K9me1/2 improved nematode tolerance to oxidative stress and heat stress. Notably, the inhibition of worm H3K9me1/2 methyltransferases by a human G9a inhibitor also extends the lifespan of *daf-2* animals. Thus, targeting H3K9me1/2 modification may be a new means for the treatment of aging and age-related diseases.

Lifespan is controlled by both genetic and epigenetic factors. The perturbation of insulin/insulin-like signaling (IIS), target of rapamycin (TOR) pathway, and mitochondrial functions have been shown to extensively modulate the aging process and lifespan. Epigenetic marks, including histone acetylation and methylation, are altered during aging and regulate lifespan in a number of species (Lee et al., 2000; Lund et al., 2002; Bennett-Baker et al., 2003; Lu et al., 2004). In *C. elegans*, both H3K4 and H3K27 methylation are involved in lifespan limitation. Animals with reduced H3K4me, for example, in *wdr-5, ash-2,* and *set-2* mutants, live longer than wild-type animals (GREER et al. 2010). Interestingly, perturbation of H3K27 methylation, via either a decrease in *mes-2* mutants or an increase in *utx-1* mutants, is associated with enhanced longevity, suggesting context-dependent lifespan regulation (17, 43).

H3K9 methylation is usually considered a repressive modification and is associated with heterochromatin. However, H3K9me2 and H3K9me3 each has distinct functions. In *C. elegans*, H3K9me2, rather than H3K9me3, is most closely associated with canonical heterochromatin factors such as HP1 (44–46). SET-25 is a known H3K9 trimethyltransferase that silences novel insertions of RNA or DNA transposons and represses tissue-specific genes during development. SET-25 is recruited to targets either by H3K9me2 deposited by MET-2 or by somatic Argonaute NRDE-3 and small RNAs (23, 47). However, we did not observe a significant change in lifespan in *set-25* mutants. Consistently, the depletion of the two heterochromatin factors HPL-1 and HPL-2 failed to alter lifespan. It was postulated that losing repressive chromatin is detrimental to lifespan (48, 49). In humans, two premature aging diseases are caused by mutations in lamins that reduce heterochromatin and disrupt its nuclear localization (50). In *C. elegans*, *Drosophila,* and mammals, heterochromatin decreases as individuals grow older (51–53). Additionally, across eukaryotes, mutations that increase repressive chromatin extend lifespan (17, 54–56). The depletion of SET-25, HPL-1 and HPL-2 failed to induce detectable lifespan alterations, suggesting that heterochromatin itself may not be sufficient to perturb lifespan.

MET-2, a mammalian H3K9 methyltransferase SETDB1 homolog (LOYOLA et al. 2006), monomethylates and dimethylates H3K9 in *C. elegans* (TOWBIN et al. 2012). MET-2 is necessary both for a normal lifespan (TIAN et al. 2016) and for the lifespan extension of *wdr-5* mutants (LEE et al. 2019). In *met-2* mutants, the lifespan is shortened. However, the lifespan was significantly extended in *daf-2;met-2* double mutants, suggesting context-dependent lifespan regulation by epigenetic machinery. SET-6 is a putative H3K9me2/3 methyltransferase that was postulated to prevent healthy aging rather than modulate the normal lifespan (22). SET-6 accelerates behavioral deterioration in *C. elegans* by reducing mitochondrial function and repressing the expression of nuclear-encoded mitochondrial proteins. Here, we found that the depletion of *set-6* dramatically increased the lifespan of *daf-2* animals, which further supports the context-dependent lifespan regulation model.

In this work, we identified a new class of lifespan-limiting histone methyltransferases, including MET-2, SET-6, SET-19, SET-20, SET-21, SET-32 and SET-33. The depletion of these proteins induced synergistic lifespan extension in *daf-2* animals. The *C. elegans* genome encodes 38 SET domain-containing proteins, most of which have no reported functions (Andersen and Horvitz 2007). Here, we showed that all these lifespan-limiting SET proteins are involved in H3K9me1/2 modification. However, further work is required to confirm their enzymatic activities. Consistent with their putative H3K9me1/2 methylation factors, treating *daf-2* animals with the human G9a inhibitor A-366 reduced H3K9me2 levels in *C. elegans* and extended the lifespan of *daf-2* animals.

Additionally, we identified a number of genes required for H3K9me1/2-involved lifespan extension. Among them, *nhr-62* is known to be required for longevity in *C. elegans* (39). SOD-3 is a superoxide dismutase that is involved in the removal of superoxide radicals. ASM-2 is an ortholog of human acid sphingomyelinase (ASM) and is involved in ceramide biosynthetic processes and sphingomyelin catabolic processes (41, 42). The *C. elegans* genome encodes three ASM homologs, *asm-1*, *asm-2* and *asm-3*. Among them, ASM-3 is most closely related to human ASM. During development and aging, ceramide and sphingosine accumulate (57). Interestingly, monounsaturated fatty acid (MUFA) accumulation is necessary for the lifespan extension of H3K4me3-methyltransferase-deficient worms, and dietary MUFAs are sufficient to extend lifespan (58). However, when *asm-1, asm-2,* or *asm-3* was inactivated by RNAi knockdown, a modest lifespan extension phenotype was observed (59). Further investigation is required to examine how and why these genes are involved in lifespan regulation.

Stichodactyla toxin (ShK, ShkT) is a 35-residue basic peptide from the sea anemone Stichodactyla helianthus that blocks a number of potassium channels with nanomolar to picomolar potency. We found that F35E8.7 and Y39G8B.7 are required for lifespan extension of H3K9me1/2-deficient mutants. Human proteins containing ShK-like domains are MMP-23 (matrix metalloprotease 23) and MFAP-2 (microfibril-associated glycoprotein 2). Whether modulating the expression of these two proteins or their downstream targets can improve longevity or healthy lifespan will stimulate much drug potential.

## Materials and methods

### Strains

Bristol strain N2 was used as the standard wild-type strain. All strains were grown at 20 °C unless specified. The strains used in this study are listed in Supplementary Table S1.

### Lifespan assay

Lifespan assays were performed at 20℃. Worm populations were synchronized by placing young adult worms on NGM plates seeded with the E. coli strain OP50-1 (unless otherwise noted) for 4–6 hours and then removed. The hatching day was counted as day one for all lifespan measurements. Worms were transferred every other day to new plates to eliminate confounding progeny. Animals were scored as alive or dead every 2 (before 52 days) or 4 (after 52 days) days. Worms were scored as dead if they did not respond to repeated prods with a platinum pick. Worms were censored if they crawled off the plate or died from vulval bursting and bagging. For each lifespan assay, 90 worms were used in 3 plates (30 worms/plate).

### Brood size

L4 hermaphrodites were singled onto plates and transferred daily as adults until embryo production ceased and the progeny numbers were scored.

### Hydrogen peroxide assay

Ten synchronized worms at day 1 of adulthood were transferred to each well, which contained 1 mL of worm S-basal buffer with various concentrations of H2O2 in a 12-well plate at 20 °C. Four hours later, 100 μL of 1 mg/mL catalase (Sigma, C9322) was added to neutralize H2O2, and the mortality of worms was scored.

### Heat-shock assay

Approximately 30 synchronized hermaphrodites at day 1 of adulthood were incubated at 35 °C, and their mortality was checked every 2 or 4 h.

### G9a inhibitor treatment

Several P0 young adult worms were placed into fresh NGM plates seeded with E. coli strain OP50 (with the same concentration of G9a inhibitors in both NGM plates and OP50 liquid). Four or five days later, several F1 young adult worms were placed into new NGM plates with OP50 and G9a inhibitor. Then, we single out the F2 worms for other assays.

### Western blot

Embryos or L4-stage worms were harvested and washed three times with M9 buffer. Samples were frozen in −80 °C. Ten minutes at 95 °C in 1X protein dye (62.5 mM Tris pH 6.8, 10% glycerol, 2% SDS, 5% β-mercaptoethanol, 0.2% bromophenol blue) was sufficient to expose worm proteins. The next step was spin for 1 minute at high speed to remove insoluble components, then quickly transfer supernatant into a new tube (on ice) and immediately run on gel or store aliquots at −80 °C. Proteins were resolved by SDS–PAGE on gradient gels (10% separation gel, 5% spacer gel) and transferred to a Hybond-ECL membrane. After washing with 1x TBST buffer (Sangon Biotech, Shanghai) and blocking with 5% milk-TBST, the membrane was incubated overnight at 4 °C with antibodies (listed below). The membrane was washed three times for 10 minutes each with 1x TBST and then incubated with secondary antibodies at room temperature for two hours. The membrane was washed three times for 10 minutes with 1x TBST and then visualized.

The primary antibodies used were β-actin (Beyotime, AF5003), H3 (Abcam, ab1791), H3K4me1 (Abcam, ab176877), H3K4me2 (Abcam, ab32356), H3K4me3 (Abcam, ab8580), H3K9me1 (Abcam, ab9045), H3K9me2 #1 (Abcam, ab1220), H3K9me2 #2 (Abcam, ab176882), H3K9me3 (Millipore, 07–523), H3K27me1 (Abcam, ab194688), H3K27me2 (Abcam, ab24684), H3K27me3 (Millipore, 07–449), H3K36me1 (Abcam, ab9048), H3K36me2 (Abcam, ab9049), H3K36me3 (Abcam, ab9050), H3K23me2 (Active Motif, 39653), and H3K23me3 (Active Motif, 61499). The secondary antibodies used were goat anti-mouse (Beyotime, A0216) and goat anti-rabbit (Abcam, ab205718) antibodies.

### Construction of deletion mutants

For gene deletions, triple sgRNA-guided chromosome deletion was conducted as previously described (60). To construct sgRNA expression vectors, the 20 bp *unc-119* sgRNA guide sequence in the pU6::unc-119 sgRNA(F+E) vector was replaced with different sgRNA guide sequences. Addgene plasmid #47549 was used to express Cas9 II protein. Plasmid mixtures containing 30 ng/µl of each of the three or four sgRNA expression vectors, 50 ng/µl Cas9 II-expressing plasmid, and 5 ng/µl pCFJ90 were coinjected into *tofu-5::gfp::3xflag (ustIS026)* animals. Deletion mutants were screened by PCR amplification and confirmed by sequencing. The sgRNA sequences are listed in Supplementary Table S2.

### RNA isolation

Synchronized L4 worms were sonicated in sonication buffer (20 mM Tris-HCl [pH 7.5], 200 mM NaCl, 2.5 mM MgCl2, and 0.5% NP40). The eluates were incubated with TRIzol reagent followed by isopropanol precipitation and DNase I digestion. mRNA was purified from total RNA using poly-T oligo-attached magnetic beads. Sequencing was performed with a HiSeqTen instrument reading 150 base paired-end reads.

### RNA-seq analysis

The Illumina-generated raw reads were first filtered to remove adaptors, low-quality tags and contaminants to obtain clean reads at Novogene. The clean reads were mapped to the reference genome of ce10 via TopHat software (version 2.1.1). Gene expression levels were determined by the fragments per kilobase of transcript per million mapped reads (FPKM).

### qRT-PCR for mRNA

Total RNA was reverse transcribed into cDNA using the GoScript Reverse Transcription System (Promega) and quantified by qPCR using SYBR GREEN mix (Vazyme Q111–02, Nanjing) with a MyIQ2 real-time PCR system. Levels of *ama-1* mRNA were used as internal controls for sample normalization. Data are expressed as fold changes relative to those of *daf-2 (e1370)* animals. The data analysis was performed using a ΔΔCT approach. The primers used for qRT–PCR are listed in Supplementary Table S3.

### ChIP-qPCR

Chromatin immunoprecipitation (ChIP) experiments were performed as previously described with L4 staged animals. After crosslinking, samples were resuspended in 1 ml FA buffer (50 mM Tris/HCl [pH 7.5], 1 mM EDTA, 1% Triton X-100, 0.1% sodium deoxycholate, and 150 mM NaCl) with a proteinase inhibitor tablet (Roche no. 05056489001) and sonicated for 20 cycles at high output (each cycle: 30 s on and 30 s off) with a Bioruptor plus. Lysates were precleared and then immunoprecipitated with 2 μl anti-histone H3 (monomethyl K9) antibody (Abcam no. ab9045), 2 μl anti-histone H3 (dimethyl K9) antibody (Abcam no. mAbcam1220) or 2 μl anti-trimethylated H3K9 antibody (Millipore no. 07–523). ChIP signals were normalized to coimmunoprecipitated *ama-1* and then expressed as fold changes relative to that of *daf-2 (e1370)* animals. qRT–PCR primers for ChIP assays are listed in Supplementary Table S4.

### Statistics

Bar graphs with error bars are presented as the mean and standard deviation. All of the experiments were conducted with independent *C. elegans* animals for the indicated N times. Statistical analysis was performed with a two-tailed Student’s t-test.

### Data availability

All raw and normalized sequencing data have been deposited in the Gene Expression Omnibus under submission number GSExxx.

## Acknowledgments

We are grateful to the members of the Guang lab for their comments. We are grateful to the International *C. elegans* Gene Knockout Consortium, and the National Bioresource Project for providing the strains. Some strains were provided by the CGC, which is funded by NIH Office of Research Infrastructure Programs (P40 OD010440). This work was supported by grants from the Strategic Priority Research Program of the Chinese Academy of Sciences (XDB39010600), the National Key R&D Program of China (2019YFA0802600, and 2018YFC1004500), the National Natural Science Foundation of China (91940303, 31870812, 32070619, 31871300 and 31900434), the China Postdoctoral Science Foundation (2018M632542, 2019T120543), and Anhui Natural Science Foundation (1808085QC82 and 1908085QC96). This study was supported, in part, by the Fundamental Research Funds for the Central Universities.

## Author Contributions

D.C., X.C., S.G. and X.F. designed the project; M.H., M.H., C.Z. performed research and data analysis. X.F. and S.G. wrote the paper.

## Supporting online materials

Figs. S1 to S7

Tables S1 to S4

## Supplementary figure legends

**Figure S1.**
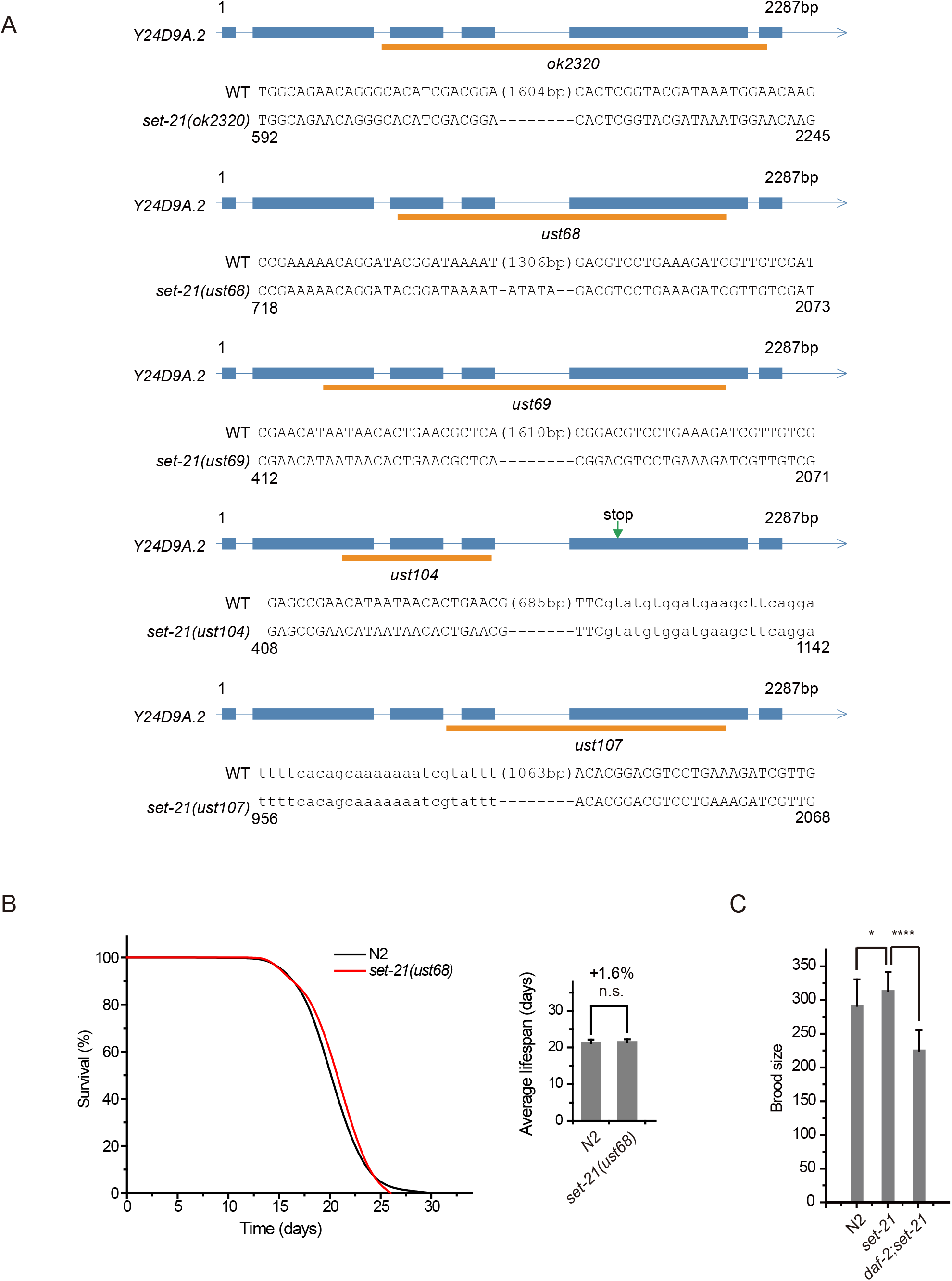
(a) Gene structure and the alleles of *set-21*. The *ust* alleles were generated by dual sgRNA-mediated CRISPR/Cas9 technology. Orange bars indicate deleted regions in gene loci. (b) Deletion of *set-21* didn’t extend the lifespan in wildtype worms, but *set-21;daf-2* double mutant revealed lifespan extension. (Left) Survival curves and (right) average lifespan of the indicated animals. The percentage of change was compared to the average lifespan of *daf-2* animals. mean ± s.e.m. of three independent experiments. ****P < 0.0001; n.s., not significant. (c) The brood size of indicated animals. Data are presented as the mean ± s.e.m. of at least 20 worms. *P < 0.05; **** P < 0.0001.

**Figure S2.**
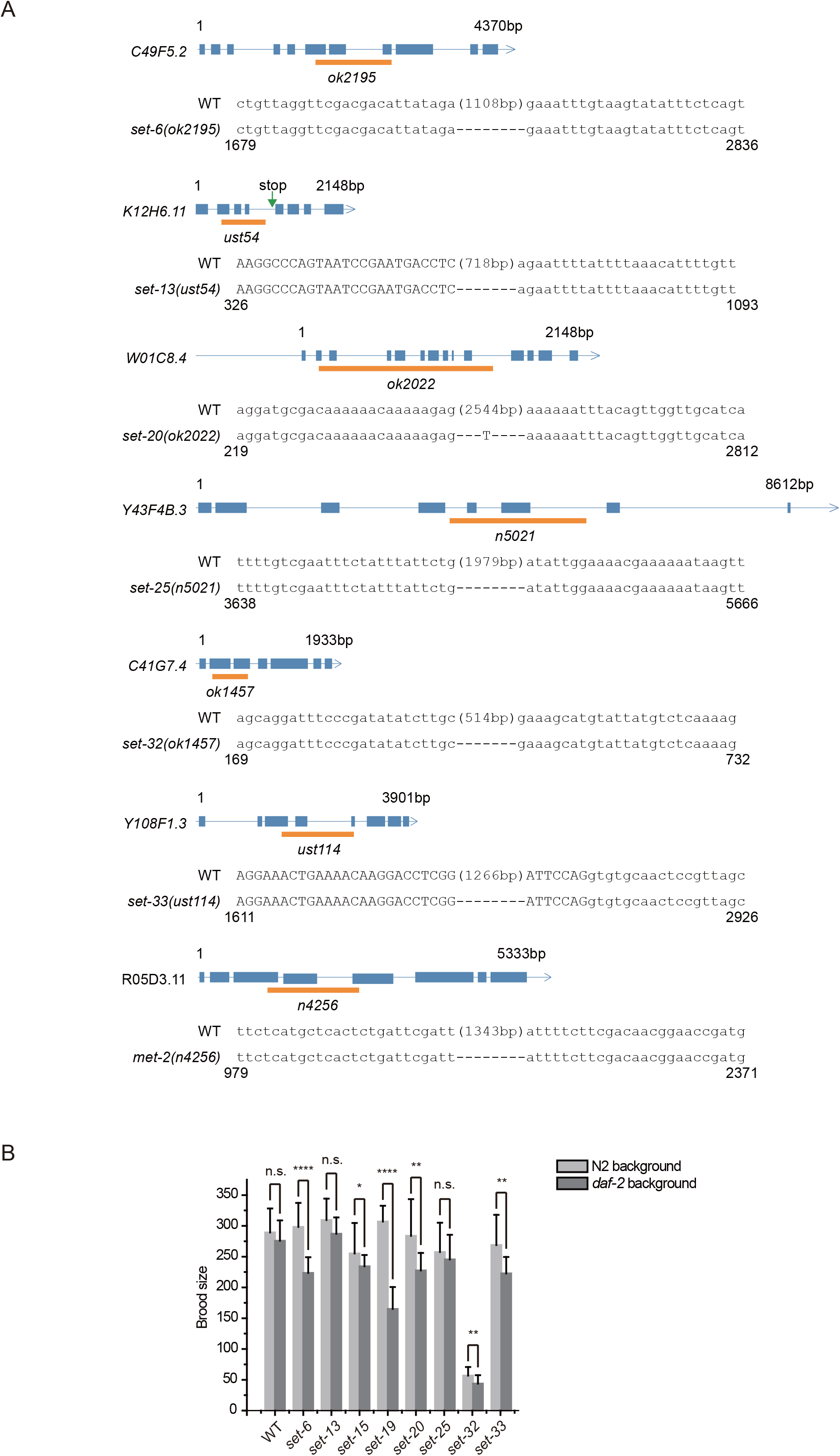
Alleles of *set* genes. (a) Gene structure and the alleles of *set* genes. The *ust* alleles were generated by dual sgRNA-mediated CRISPR/Cas9 technology. Orange bars indicate deleted regions in gene loci. (b) Brood size of indicated animals. Data are presented as the mean ± s.e.m. of at least 20 worms. *P < 0.05; **P < 0.01; ***P < 0.001; ****P < 0.0001; n.s., not significant.

**Figure S3.**
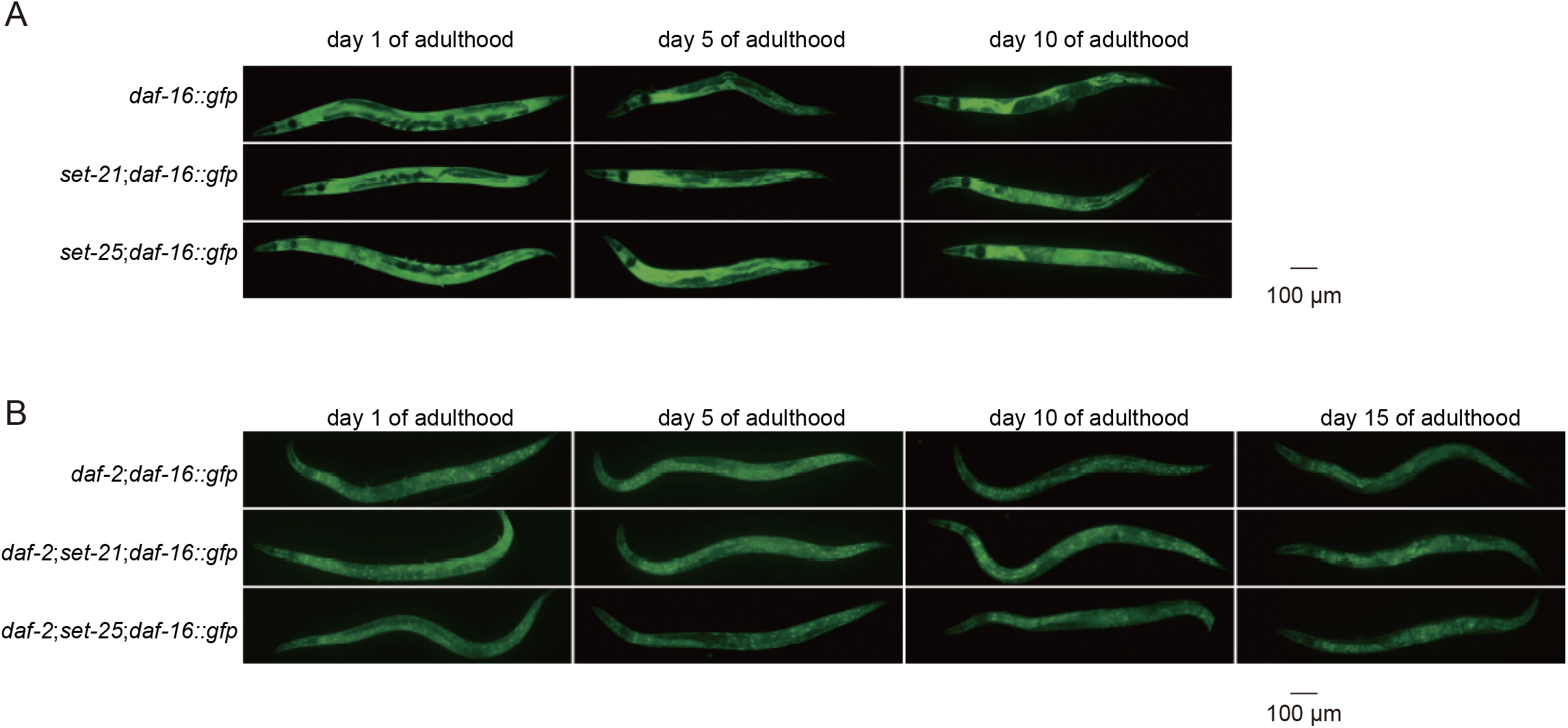
Fluorescent images of GFP::DAF-16 in the indicated young adult animals.

**Figure S4.**
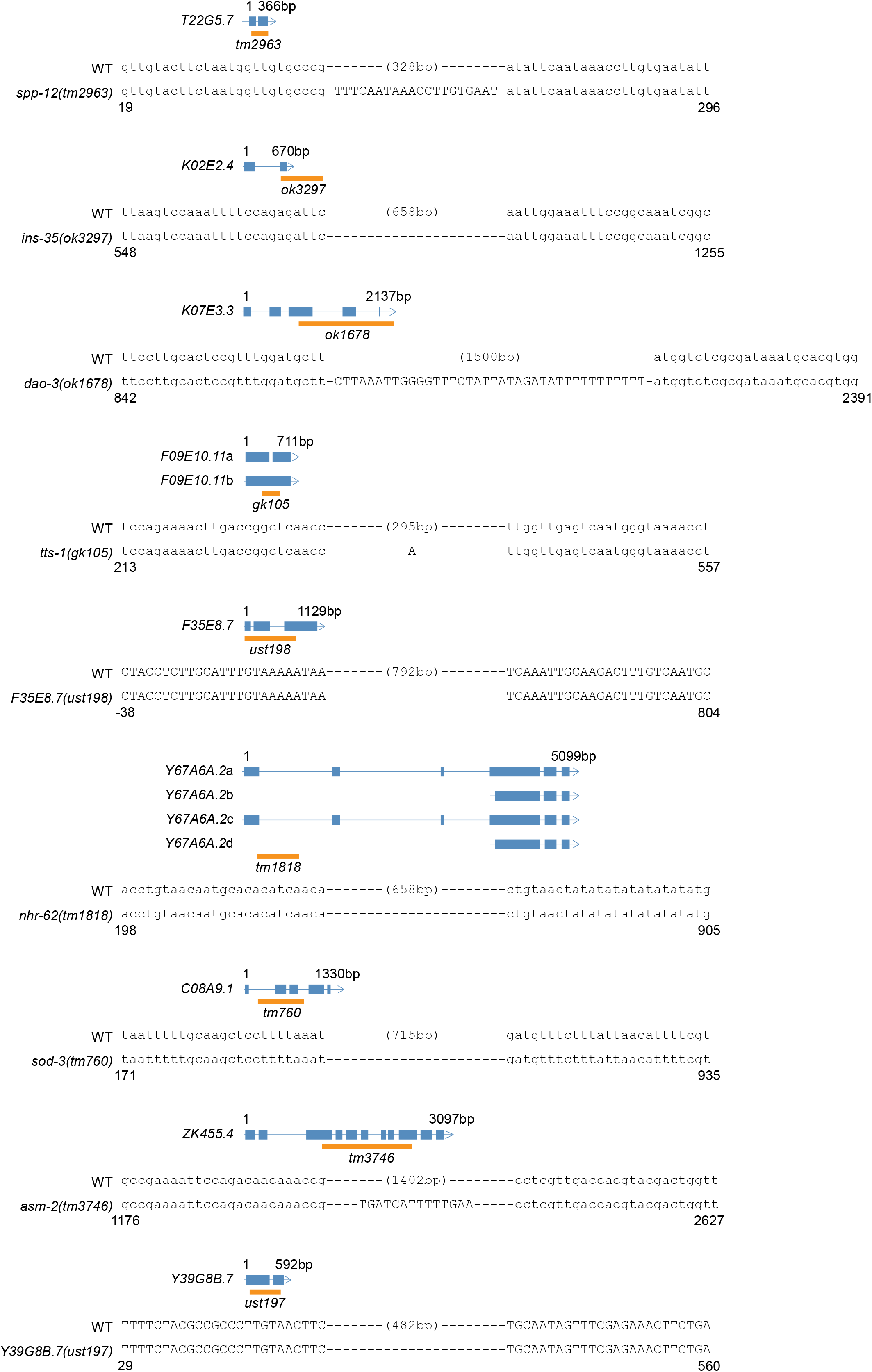
Gene structure and the alleles of indicated genes. The *ust* alleles were generated by dual sgRNA-mediated CRISPR/Cas9 technology. Orange bars indicate deleted regions in gene loci.

**Figure S5.**
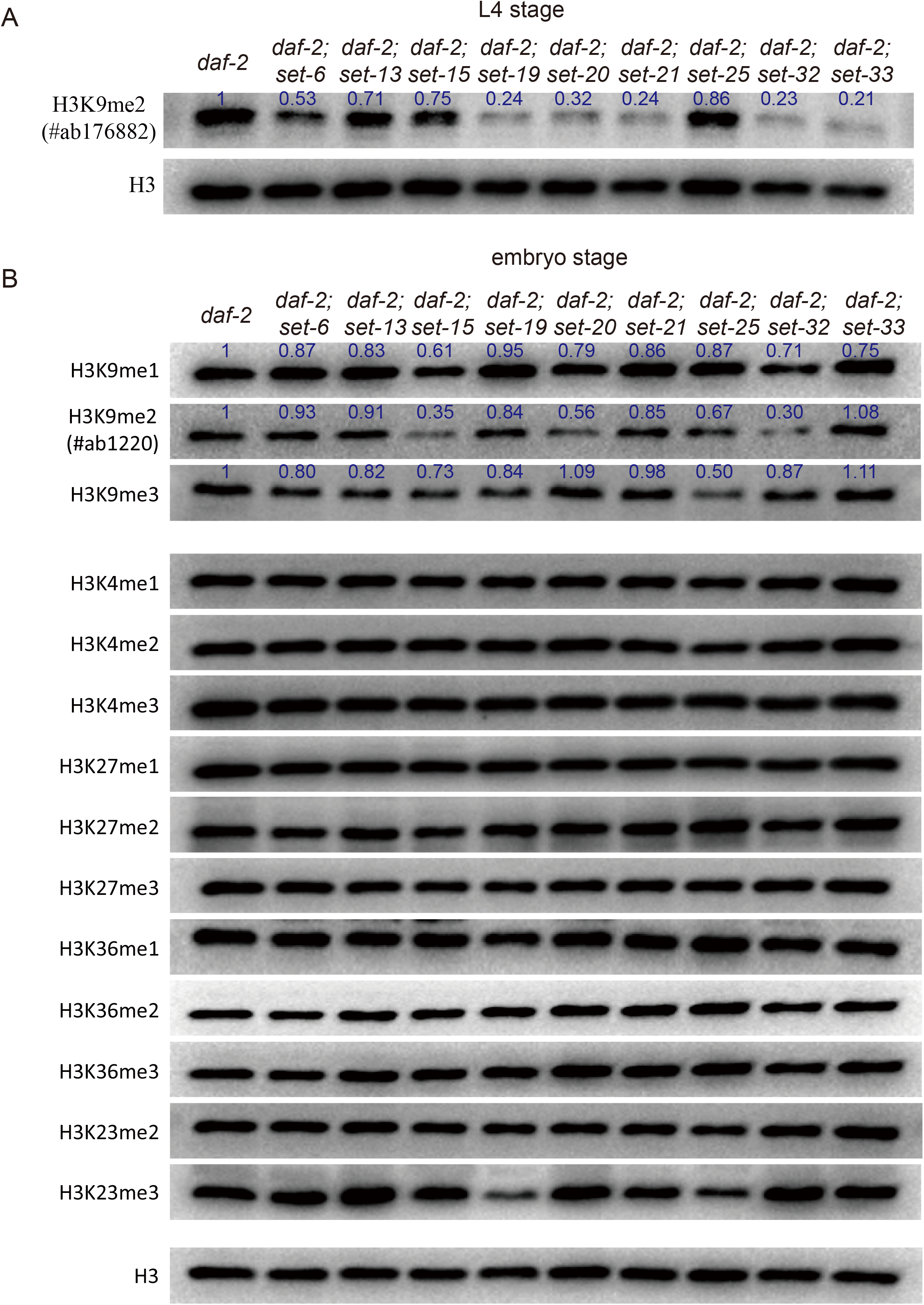
Western blotting of (a) L4 and (b) embryos with the indicated antibodies. Numbers in the picture indicate the brightness of bands measured by ImageJ. The original files of the full raw unedited blots and figures with the uncropped blots with the relevant bands clearly labelled are provided in the source data 3.

**Figure S6.**
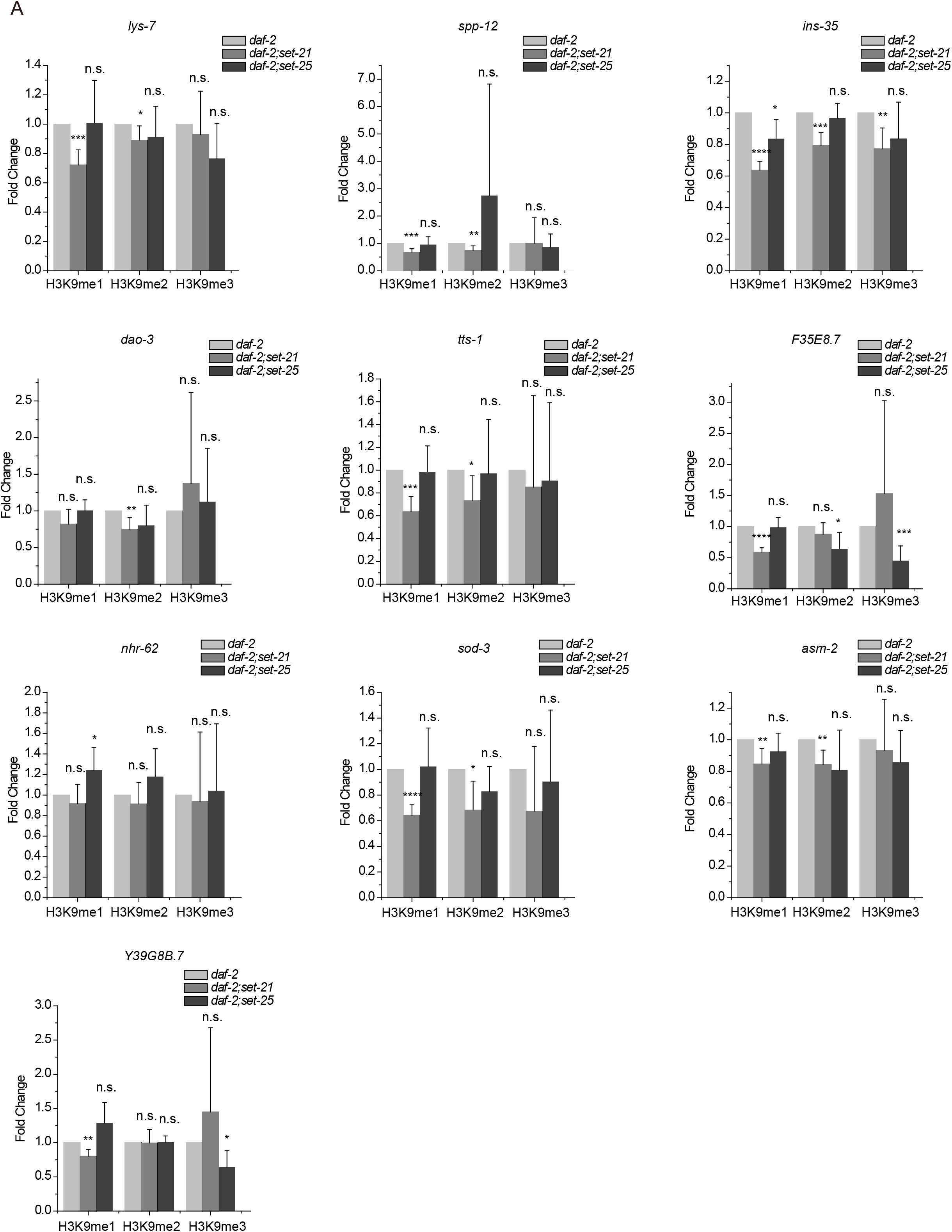
Chromatin immunoprecipitation of histone methylation marks in the indicated genes at the L4 stage. Data are presented as ratios of H3K9 methylation levels in *daf-2;set-21* and *daf-2;set-25* versus daf-2 animals. H3K9 signals from *ama-1* were used as an internal control for ChIP normalization. Data are presented as the mean ± s.e.m. of five independent experiments. *P < 0.05; **P < 0.01; ***P < 0.001; ****P < 0.0001; n.s., not significant.

**Figure S7.**
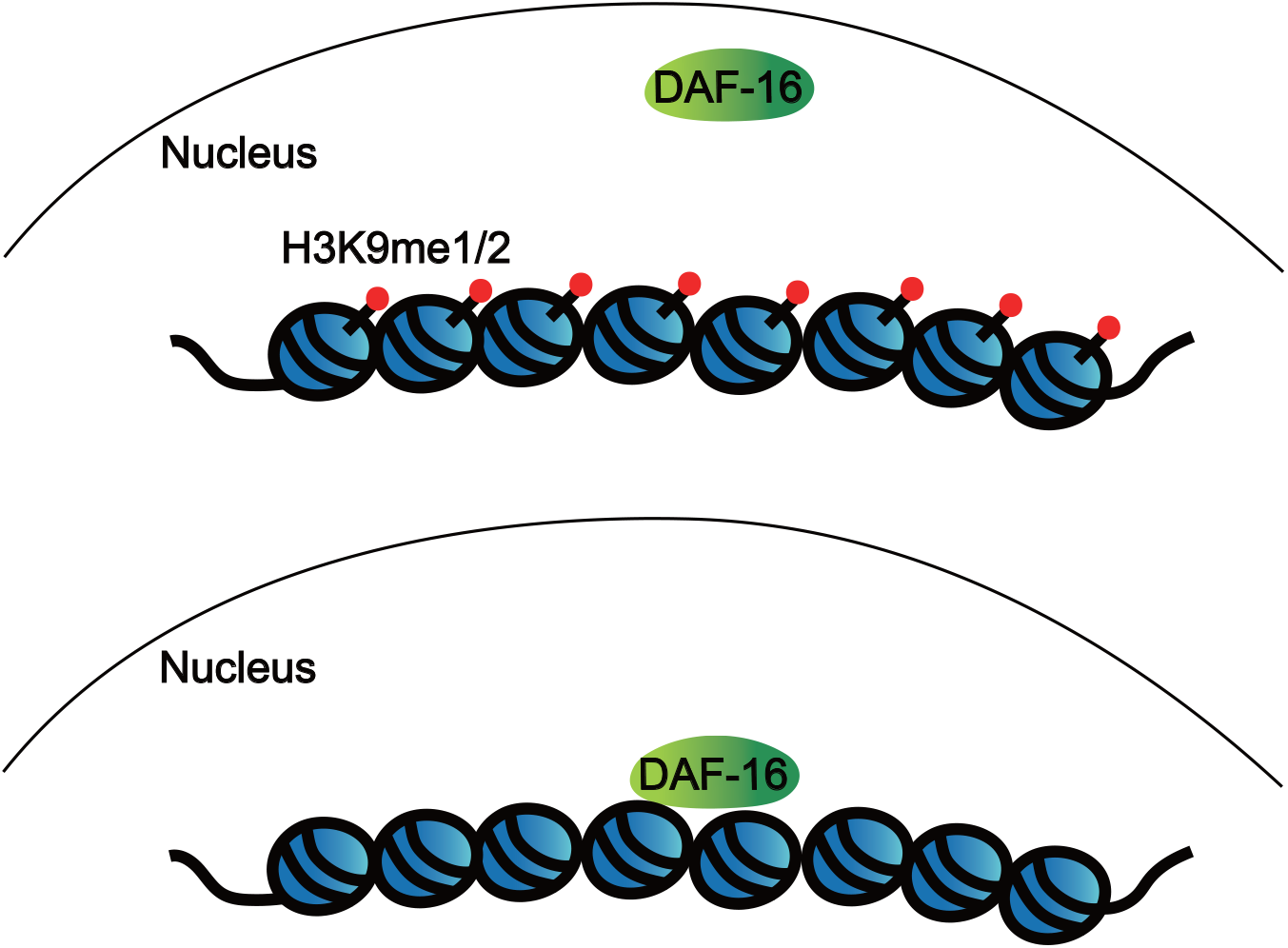
A working model of H3K9me1/2 marks regulating lifespan of *C. elegans* via modulating the association of DAF-16 to targeted genes. The loss of H3K9me1/2 increases the binding of DAF-16 to Class I genes and promotes the expression of lifespan-promoting and anti-stress genes.

**Table S1.** List of strains used in this study.

| strain | genotype |
| --- | --- |
| SHG463 | *set-13(ust54)* II |
| SHG563 | *daf-2(e1370)* III; *set-21(ok2320)* IV |
| SHG564 | *daf-2(e1370)* III; *set-13(ust54)* II |
| SHG565 | *daf-2(e1370)* III; *daf-16(mu86)* I |
| SHG728 | *set-6(ok2195)* X; *daf-2(e1370)* III |
| SHG729 | *set-19(ok1813)* X; *daf-2(e1370)* III |
| SHG730 | *set-20(ok2022)* X; *daf-2(e1370)* III |
| SHG731 | *set-25(n5021)* III; *daf-2(e1370)* III |
| SHG732 | *set-32(ok1457)* I; *daf-2(e1370)* III |
| SHG733 | *rsks-1(ok1255)* III; *set-21(ust68)* IV |
| SHG734 | *rsks-1(ok1255)* III; *daf-2(e1370)* III; *set-21(ust68)* IV |
| SHG792 | *daf-2(e1370)* III; *daf-16(mu86)* I; *set-21(ust68)* IV |
| SHG822 | *set-21(ust68)* IV; *zIs356 [daf-16p::daf-16a/b::GFP + rol-6(su1006)]* IV |
| SHG841 | *daf-2(e1370)* III; *hpl-2(tm1489)* III |
| SHG846 | *daf-2(e1370)* III; *set-21(ust68)* IV; *set-6(ok2195)* X |
| SHG847 | *daf-2(e1370)* III; *set-21(ust68)* IV; *set-19(ok1813)* X |
| SHG848 | *daf-2(e1370)* III; *set-21(ust68)* IV; *set-20(ok2022)* X |
| SHG849 | *daf-2(e1370)* III; *set-21(ust68)* IV; *set-32(ok1457)* I |
| SHG858 | *set-21(ust104)* IV |
| SHG861 | *set-21(ust107)* IV |
| SHG869 | *set-33(ust114)* X |
| SHG1131 | *rsks-1(ok1255)* III; *daf-2(e1370)* III |
| SHG1133 | *daf-2(e1370)* III |
| SHG1136 | *daf-16(mu86)* I; *set-21(ust69)* IV |
| SHG1137 | *daf-2(e1370)* III; *set-21(ust69)* IV |
| SHG1138 | *set-21(ust68)* IV |
| SHG1139 | *set-21(ust69)* IV |
| SHG1140 | *daf-2(e1370)* III; *set-21(ust68)* IV |
| SHG1141 | *daf-16(mu86)* I; *set-21(ust68)* IV |
| SHG1145 | *daf-2(e1370)* III; *set-15(ok3291)* IV |
| SHG1146 | *daf-2(e1370)* III; *set-21(ust68)* IV; *set-2(ok952)* III |
| SHG1147 | *daf-2(e1370)* III; *set-2(ok952)* III |
| SHG1148 | *daf-2(e1370)* III; *hpl-1(tm1624)* X |
| SHG1149 | *daf-16(mu86)* I; *daf-2(e1370)* III; *set-21(ust68)* IV |
| SHG1150 | *daf-2(e1370)* III; *set-21(ust104)* IV |
| SHG1151 | *daf-2(e1370)* III; *set-33(ust114)* X |
| SHG1164 | *daf-2(e1370)* III; *set-21(ust68)* IV; *asm-2(tm3746)* X |
| SHG1165 | *daf-2(e1370)* III; *set-21(ust68)* IV; *ins-35(ok3297)* V |
| SHG1166 | *daf-2(e1370)* III; *set-21(ust68)* IV; *lys-7(ok1384)* V |
| SHG1167 | *daf-2(e1370)* III; *set-21(ust68)* IV; *nhr-62(tm1818)* I |
| SHG1168 | *daf-2(e1370)* III; *set-21(ust68)* IV; *spp-12(tm2963)* V |
| SHG1169 | *daf-2(e1370)* III; *set-21(ust68)* IV; *tts-1(gk105)* X |
| SHG1170 | *daf-2(e1370)* III; *set-21(ust68)* IV; *sod-3(tm760)* X |
| SHG1171 | *daf-2(e1370)* III; *set-21(ust68)* IV; *dao-3(ok1678)* X |
| SHG1172 | *daf-2(e1370)* III; *set-21(ust68)* IV; *F35E8.7(ust198)* V |
| SHG1173 | *daf-2(e1370)* III; *set-21(ust68)* IV; *Y39G8B.7(ust197)* II |
| SHG1175 | *daf-16(mu86)* I; *set-2(ok952)* III |
| SHG1177 | *daf-2(e1370)* III; *set-21(ust68)* IV; *set-33(ust114)* X |
| SHG1178 | *daf-2(e1370)* III; *set-21(ust106)* IV |
| SHG1179 | *daf-2(e1370)* III; *set-21(ust107)* IV |
| SHG1197 | *met-2(n4256)* III |
| SHG1199 | *met-2(n4256)* III; *daf-2(e1370)* III |
| SHG1367 | *daf-2(e1370)* III; *zIs356 [daf-16p::daf-16a/b::GFP + rol-6(su1006)]* IV |
| SHG1368 | *daf-2(e1370)* III; *set-25(n5021)* III;<br>*zIs356 [daf-16p::daf-16a/b::GFP + rol-6(su1006)]* IV |
| SHG1369 | *daf-2(e1370)* III; *set-21(ust68)* IV;<br>*zIs356 [daf-16p::daf-16a/b::GFP + rol-6(su1006)]* IV |

**Table S2.**
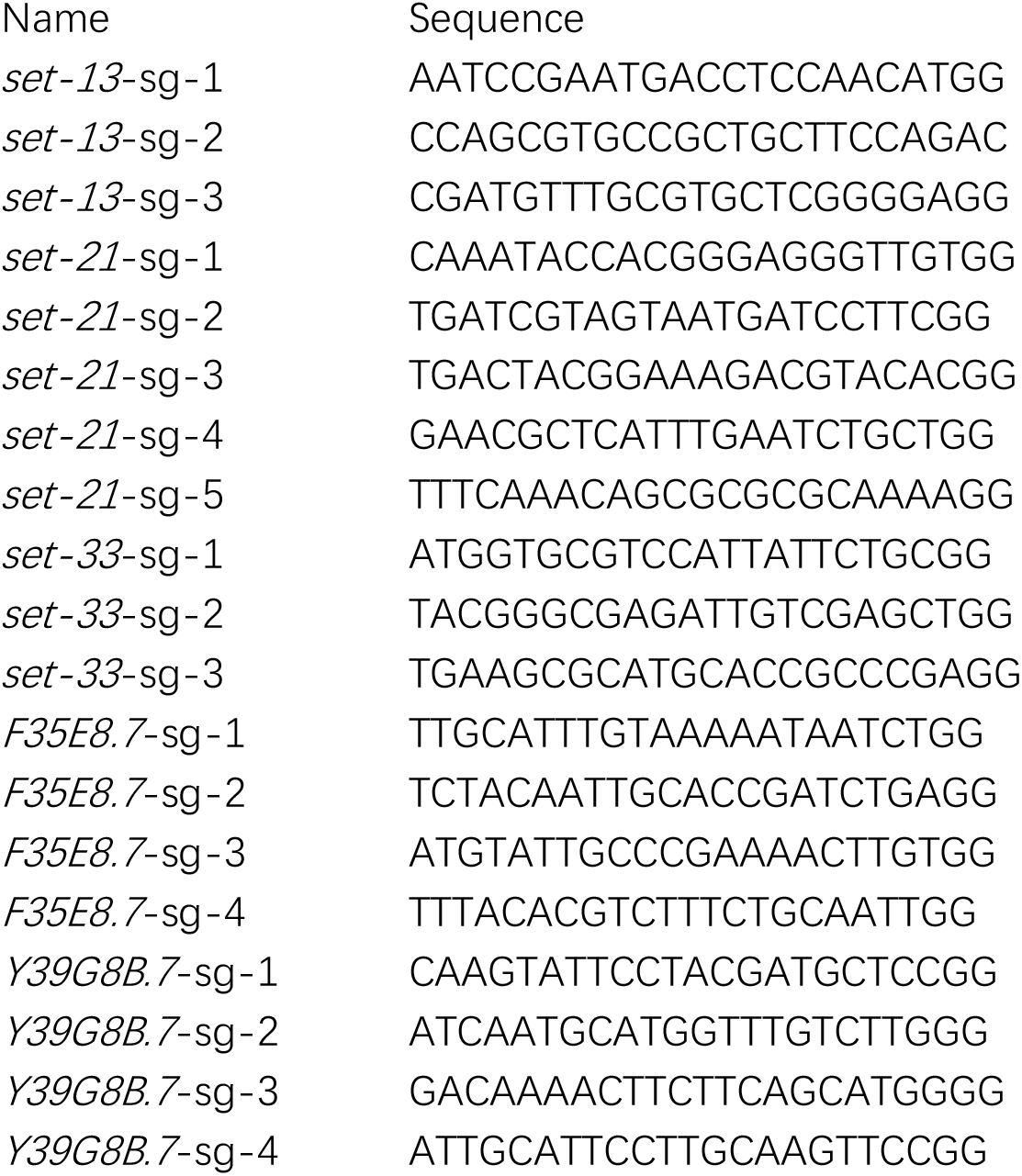
sgRNA sequences for CRISPR/Cas9-directed gene editing technology.

**Table S3.**
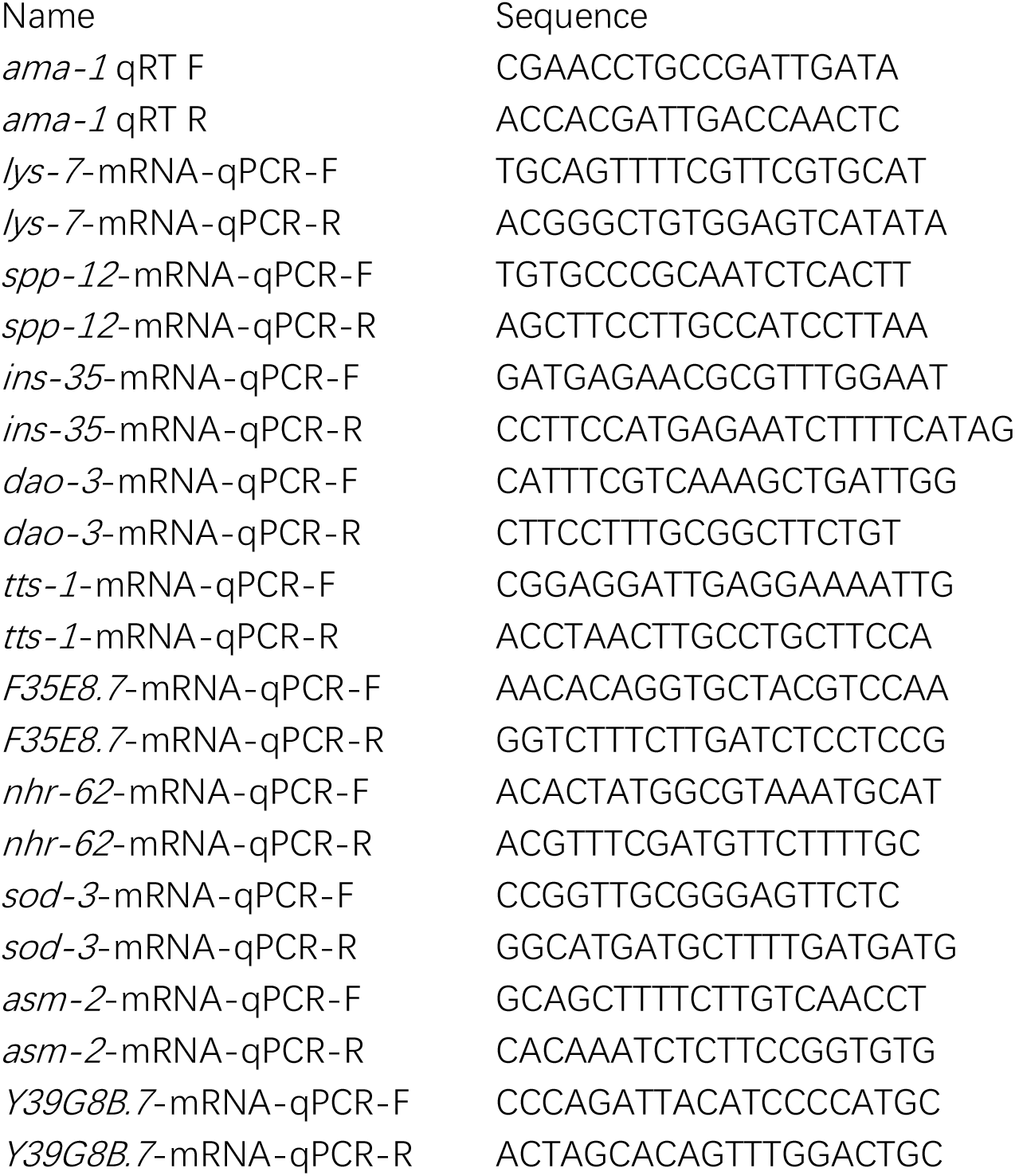
List of primers used in mRNA qPCR.

**Table S4.**
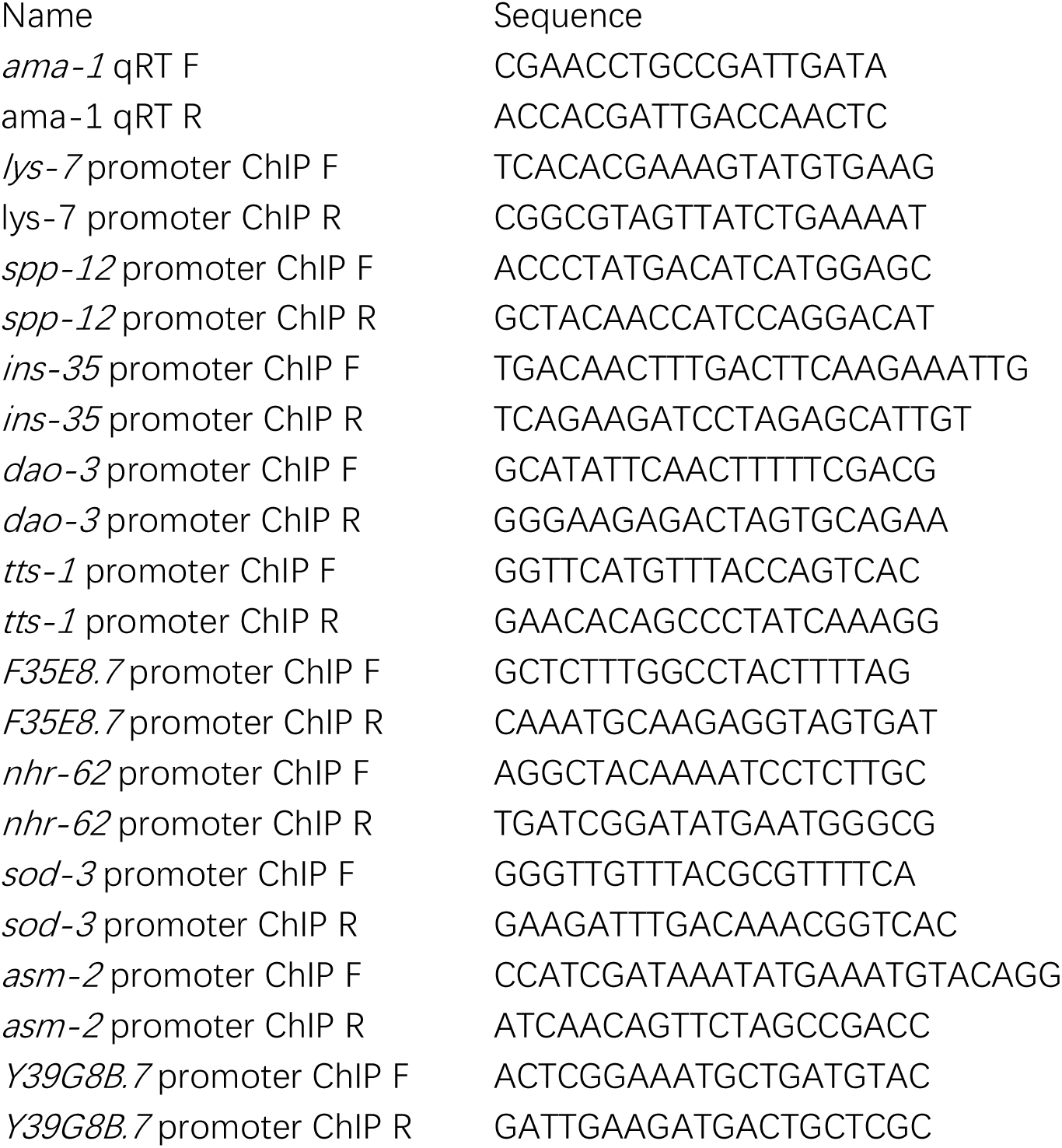
List of primers used in ChIP-qPCR.

